# Impact of iron-enriched diets on the nutritional composition and microbiome diversity of house crickets (*Acheta domesticus*)

**DOI:** 10.64898/2026.11.06.764535

**Authors:** M.J. Mayen, C. Schüßler, S.O. Adejo, G.O. Obochi, H. Haase, M. Maares, S. Mbedi, S. Sparmann, O.K. Schlüter, C. Rafaluk, C. Keil, G. Rossi

## Abstract

Iron deficiency (ID) and iron-deficiency anaemia (IDA) pose significant global health challenges. ID and IDA are particularly widespread in Sub-Saharan Africa and South Asian countries, where reliance on plant-based foods is high. Edible insects provide an alternative nutritious food source, rich in both protein and essential minerals and trace elements. Little is known about how dietary iron manipulation affects the trace element bioaccumulation and microbiome community composition of edible insects. This study examined the impact of dietary iron supplementation on the growth, survival, nutrient profile, and microbiome of *Acheta domesticus*. Experimental diets were formulated using iron (II) sulphate heptahydrate, resulting in final iron concentrations ranging from 333.45 mg Fe/kg (control diet) to 6923.39 mg Fe/kg (100 mM Fe diet). Iron supplementation significantly altered the cricket trace element profile, with iron levels increasing fivefold from basal 21.38 mg/kg to 118.34 mg/kg in the highest iron treatment group. Concurrently, manganese levels showed a strong negative correlation with dietary iron, while zinc and cadmium displayed non-significant downward trends. High-iron diets statistically altered dry matter, chitin, and TBARS, whereas carbohydrates, protein, glycogen, and lipids remained stable. Amino acid profiles shifted progressively, with branched-chain, sulphur-containing, and aromatic amino acids peaking in the 25 mM Fe group. In contrast, the diversity and overall structure of the microbial community remained unaffected across all treatments. In conclusion, tailoring dietary iron can significantly boost the iron content of *Acheta domesticus* without compromising host survival or altering core microbial structures. To optimize insect rearing strategies aimed at alleviating human micronutrient deficiencies, a feeding dose of 10–25 mmol Fe/kg is recommended.

## Introduction

Iron deficiency is among the most prevalent micronutrient deficiencies globally and a major contributor to the global burden of disease (Lee *et al*., 2025). The World Health Organization (WHO) estimated that over 2 billion people are affected by iron deficiency (ID) and iron deficiency anaemia (IDA) globally. The problem is particularly profound in sub-Saharan Africa and South Asia (Pan *et al*., 2026). ID has a significant negative impact on the cognitive development of children, physical working capacity for adults, and poor birth outcomes for pregnant women (Georgieff, 2020; Pivina *et al*., 2019). Although numerous supplementation and fortification programs have been launched to improve the situation (Bouis, 2023; Kaur *et al*., 2022; Osendarp *et al*., 2018), the burden of iron deficiency persists (Bathla and Arora, 2022; Lee *et al*., 2025).

In Sub-Saharan Africa and South Asia, the ID challenge is exacerbated by the reliance on plant-based diets, which are particularly rich in non-heme iron species, which are poorly absorbed by the human body (Rousseau *et al*., 2020). Phytates, polyphenols, and calcium in these diets act as inhibitors, reducing intestinal iron absorption and bioavailability and rendering the nutritional effectiveness of plant-based diets limited (Punetha *et al*., 2026). To achieve a better iron supply, the dietary integration of non-plant food that are inherently present and culturally accepted across regions of the Global South presents a promising option (Adesogan *et al*., 2026).

Edible insects have emerged as a sustainable food source that provide adequate amounts of energy, protein, fatty acids, and micronutrients (van Huis *et al*., 2021). Of the widely consumed edible insects (Omuse et al 2024), crickets including *Acheta domesticus* (*A. domesticus*) and *Gryllus bimaculatus* have received particular attention due to their high nutritional content, characterized by high concentrations of essential micronutrients, protein, lipids, and essential amino acids (Yadav *et al*., 2026). Additionally, cricket farming is environmentally sustainable due to a low feed conversion ratio, lower greenhouse gas emissions, and lesser land and water usage than conventional livestock (Madau *et al*., 2020; van Huis and Oonincx, 2017). These attributes make crickets a highly viable option for addressing nutritional challenges (Choi *et al*., 2025; Rouco *et al*., 2026). Importantly, the nutritional composition of edible insects, including crickets, is variable and can be optimized through targeted feeding strategies. Studies have shown that dietary manipulation can modulate nutrient content, such as fatty acids, minerals/trace elements, and vitamins in edible insects (Bawa *et al*., 2020; Keil *et al*., 2020; Muzzatti *et al*., 2024). This plasticity opens avenues for biofortification, allowing specific nutrients to be enhanced through controlled feeding (Van Der Straeten *et al*., 2020). By changing the diet composition, the micronutrient content of crickets can be enhanced for human consumption and animal feed (Bawa *et al*., 2020). Applying this strategy to iron enrichment is of particular interest as the development of iron fortified edible insect could provide a potential solution to ID and IDA in the global south.

Recent human stable-isotope trials evaluating iron-fortified house crickets incorporated into food matrices demonstrated that fractional iron absorption remains rather low, typically ranging between 3% and 5% (Mwangi *et al*., 2022). In this case, however, the cricket iron enrichment achieved was only moderate; due to the technical and financial constraints of utilizing the rare stable ^57^Fe-isotopes for insect labelling (the maximum dosage offered in the feed was approximately 1.05 mmol ^57^Fe/kg diet). To elucidate the underlying mechanisms behind this low in vivo uptake, complementary *in vitro* digestion and cellular models have been employed. These laboratory models corroborate that while the chemical solubility of insect iron is high, its cellular bioavailability is severely hindered by the inhibitory properties of chitin, endogenous phytates, and mineral competition (Mwangi *et al*., 2022). One promising approach to counteract this low human absorption is to maximize the iron concentration within the cricket biomass through advanced feeding regimes—a strategy that has already proven successful in *Tenebrio molitor* (*T.* molitor) and *Hermetia illucens* (*H. illucens*) (First *et al*., 2024; First *et al*., 2026).

A key challenge for targeted iron enrichment is that high supplementation levels can trigger physiological trade-offs. Excessive iron induces oxidative stress via reactive oxygen species, restricting insect growth and overall fitness (Gomes *et al*., 2023; Gorman, 2023; Whiten *et al*., 2017). Excessive iron feeding can also trigger trace mineral dyshomeostasis; for instance, an increase in dietary iron concentration in feeding substrate has been shown to cause a significant decrease in zinc content in farmed insect species, suggesting competitive absorption pathways (First *et al*., 2024). Overfeeding insects with excessive iron poses an additional threat to their gut microbiome. This is a phenomenon known from conventional livestock production, where excessive oral iron saturates absorption, leaving unabsorbed luminal iron to fuel opportunistic pathogens, resulting in intestinal inflammation, compromising barrier integrity, and decreasing growth performance (Chen *et al*., 2020; Li *et al*., 2016; Liu *et al*., 2024). Indeed, opportunistic pathogens such as *Staphylococcus aureus* and *Pseudomonas aeruginosa* thrive in iron-rich environments (An *et al*., 2026; Ledala *et al*., 2014) and at least closely related pathogens to these are present in the insect gut (Maciel-Vergara *et al*., 2018; Nayduch *et al*., 2013; Philbert, 2025). Iron induced disruption in targeted insects could severely jeopardize host fitness, as their gut microbiome fundamentally supports digestion, nutrient uptake, immune function, and pathogen resistance, as well as posing risks to mass reared cultures and end consumers. While the direct consequences of such dietary iron overloading on farmed insects remain largely unexplored, evidence from other arthropod models confirms that maintaining intestinal iron homeostasis is equally vital for preserving microbial balance, immunity, and host fitness (Cardoso-Jaime *et al*., 2022; Hrdina and Iatsenko, 2022). These potential outcomes should be monitored and evaluated when designing enrichment protocols.

In this study, we address these literature gaps by investigating the impact of iron-enriched diets on the macronutrient and mineral/trace-element composition, as well as on the microbiome diversity and composition, of *A. domesticus* kept under controlled laboratory conditions. We hypothesized that dietary iron enrichment would increase iron accumulation in crickets and alter the gut microbiome composition. We aimed to (i) quantify the nutritional composition and total iron content of crickets fed with basal control and iron-enriched diets, (ii) evaluate shifts in the wider essential mineral and trace element profile, while assessing the accumulation of potentially toxic elements, to understand the elemental homeostasis and physiological regulation in *A. domesticus*, and (iii) characterize the microbiome diversity and community dynamics using 16S rRNA sequencing.

## 2. Materials and Methods

### 2.1 Cricket rearing

#### Diet preparation

Iron (II) sulphate heptahydrate (FeSO_4_·7 H_2_O, CAS No. 7782-63-0, purity ≥ 99.0%) was obtained from Merck (Darmstadt, Germany). For diet supplementation, aqueous solutions of FeSO_4_·7 H_2_O with varying iron concentrations were prepared (Table 1). A volume of 700 ml of each solution was thoroughly mixed with 3.5 kg of chicken feed (product number 151012; onOvo Pet Supplies GmbH, Münden, Germany) and mixed thoroughly to produce 5 diets with increasing iron content. For the control (CTR) diet, 3.5 kg of feed was mixed with 700 ml of double-distilled water. The moist feeding material was freeze-dried in two rounds (Sublimator 15, Zirbus Technology GmbH, Germany) and milled to a final size of 0.75 mm (MICROCUT MCH 20K, Stephan Machinery GmbH), considered to be optimal for cricket consumption (Van Peer *et al*., 2024). At the start of the experiment, 10 g of each feed was sampled, and its elemental composition was determined via ICP-MS according to the protocol in Section 2.2. The actual iron concentrations in the final diets are summarized in Table 1.

**Table 1.** Experimental diets used for the cricket rearing experiment.

| **Diet** | **Feed composition** | **Fe concentration in final diet (mg/kg DM)<sup>1</sup>** |
|---|---|---|
| **CTR** | 3.5 kg chicken feed + 700 mL water | 333.45 ± 12.61 |
| **Fe_1mM** | 3.5 kg chicken feed + 700 mL 5 mM FeSO<sub>4</sub> solution<br>(1.39 g FeSO<sub>4</sub>·7 H<sub>2</sub>O in 1000 mL water) | 350.31 ± 40.94 |
| **Fe_5mM** | 3.5 kg chicken feed + 700 mL 25 mM FeSO<sub>4</sub> solution<br>(6.96 g FeSO<sub>4</sub>·7 H<sub>2</sub>O in 1000 mL water) | 525.79 ± 44.94 |
| **Fe_10mM** | 3.5 kg chicken feed + 700 mL 50 mM FeSO<sub>4</sub> solution<br>(13.90 g FeSO<sub>4</sub>·7 H<sub>2</sub>O in 1000 mL water) | 806.23 ± 65.70 |
| **Fe_25mM** | 3.5 kg chicken feed + 700 mL 125 mM FeSO<sub>4</sub> solution (34.75 g FeSO<sub>4</sub>·7 H<sub>2</sub>O in 1000 mL water) | 1,668.70 ± 62.86 |
| **Fe_100mM** | 3.5 kg chicken feed + 700 mL 500 mM FeSO<sub>4</sub> solution<br>(139.01 g FeSO<sub>4</sub>·7 H<sub>2</sub>O in 1000 mL water) | 6,923.39 ± 177.41 |

<sup>1</sup> value are expressed as average ± standard error (n = 5).

#### Cricket rearing

The feeding trial was carried out at the Leibniz-Institut für Agrartechnik und Bioökonomie e.V. (ATB), Potsdam (Supplementary Figure S1). Last-stage cricket nymphs were obtained from Fauna Topics GmbH (Marback am Neckar, Germany) and placed in a transparent polypropylene box (22 L, 39x28x28 cm, Ikea Deutschland GmbH, art. 694.408.22) equipped with a lid with a hole (27x21 cm) covered by a stainless steel 1.4x1.4 mm mesh (Gazeshop.de, Braunschweig, Germany). Half egg cartons were placed in the boxes in order to provide safe hiding places for the crickets and to increase the surface area. Tap water was provided via a 100 mL water dispenser designed for birds (Nobby Pet Shop GmbH, Bocholt, Germany), filled with paper tissue to prevent crickets from drowning. Approximately 10 g of milled (Ø = 0.75 mm) chicken feed (Versele Laga NV, Deinze, Belgium) was provided in a petri dish

The rearing boxes were placed inside a climatic chamber (PolyKlima GmbH, Freising, Germany) set at 31 ± 1 °C, 50 ± 5% relative humidity, and with a photoperiod of 12:12 (L:D). Feed and water were replaced three times/week. At every feeding event, rearing boxes were cleaned with a cordless handheld vacuum cleaner (EDEKA ZENTRALE Stiftung & Co. KG, Hamburg, Germany) to remove frass and dirt.

Nymphs (approximately 100/box) were left to develop into adults in these containers for 7 days. Approximately one week after the first adult appeared, an oviposition substrate, consisting of a mixture of sand/coconut fiber (20:1) frequently sprayed with tap water (aiming to keep a moisture of 30-35% (Van Peer *et al*., 2024)), was provided in a white plastic box (10.5x10.5x3.0 cm) covered with a 1.4x1.4 mm stainless steel mesh (Gazeshop.de, Braunschweig, Germany). During the adult stage, the oviposition substrate was replaced three times, each kept for 5 days. After removing the oviposition substrate, the adults were killed by freezing, and the boxes were cleaned with tap water and soap and disinfected with 96% ethanol. The oviposition substrate was sprayed with tap water, transferred to a new clean rearing box, and incubated in the same rearing chamber used for adult rearing. Cricket eggs hatched in approximately 10 days. When the first newly hatched nymphs appeared, the mesh of the oviposition box was removed, and the oviposition substrate was spread inside the rearing box to facilitate the hatching procedure. When calculating the age of the newly hatched crickets, day 10 following the day of the introduction of the oviposition tray in the adult rearing box was taken as the starting reference (day 0). When the nymphs started to grow, water was provided in the form of a sodium polyacrylate-based moistened hydrogel (Wassergel Granulat, Tropic Shop, Nordhorn, Germany) placed inside a white plastic 50 mL hexagonal tray (Carl Roth GmbH + Co. KG, Karlsruhe, Germany), opened on one side to facilitate free access to the neonates. Milled chicken feed (Ø = 0.75 mm, Versele Laga NV, Deinze, Belgium), bio-quality carrot slices (REWE Markt GmbH, Potsdam, Germany), and sodium polyacrylate-based moistened hydrogel (Tropic Shop, Nordhorn, Germany) were provided *ad libitum*.

After 20 days of development, the nymphs were randomly divided into 6 treatment groups, with five independent replicates/treatment (5 biological replicates). Each group was reared on a different diet at defined iron concentration (Table 1). For each experimental unit, a 22-L transparent polypropylene box was prepared. Fifty 20-day-old crickets were placed in each box, and the boxes were randomly placed inside a walk-in climate room set at temperature 31 ± 1°C, relative humidity 50 ± 5%, and photoperiod 12:12 (L:D). To track the insect development, 10 randomly selected crickets from each unit were weighed every second day, and the number of escaped and dead crickets were recorded. In each box, 10 g of the corresponding diet and water as hydrogel were replaced 3 times/week (Monday, Wednesday, Friday). The rearing experiment was concluded when the first adult emerged in each specific box. This resulted in rearing being terminated differently for each group/replicate. At this stage, crickets were starved for 48 hours under original rearing conditions. Hydrogel was still provided for hydration. Harvested crickets were frozen at -80°C for further analysis.

To estimate the rearing performances of crickets, several parameters were calculated (Psarianos *et al*., 2024; Van Peer *et al*., 2024):

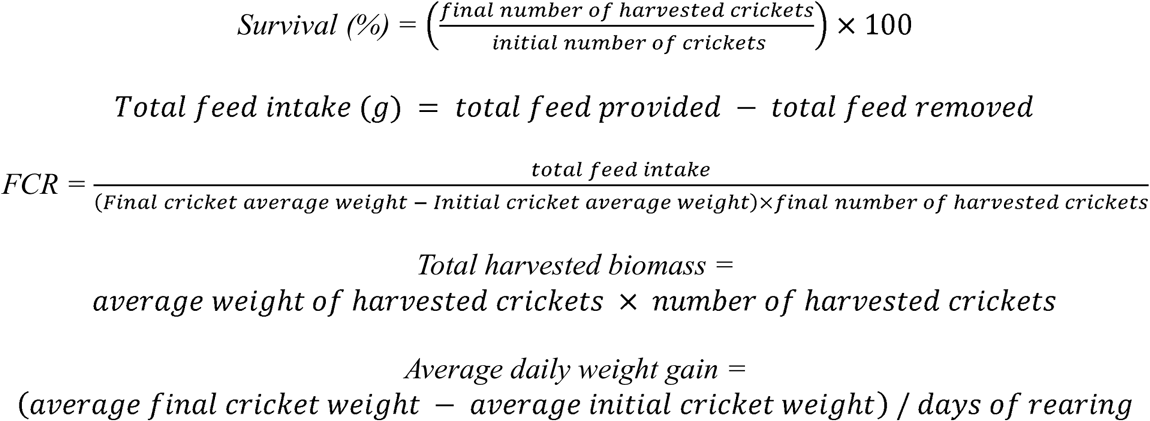

### 2.2 Chemical analyses

The insect biomass collected from each box was homogenised using a Qiagen TissueLyser II QIAGEN (Hilden, Germany) for 10 minutes at 25 Hz frequency and separated into four aliquots used for either (i) proximate analysis and amino acid profiling, (ii) mineral profiling, (iii) oxidative status (TBARS assay), (iv) microbial DNA extraction and sequencing.

#### Proximate analysis

Proximate analysis of the cricket biomass was carried out according to Cuff *et al*. (2021) with modifications. In brief, the insect biomass was dried in a forced-air electric oven (Heraeus UT_20, Thermo Fisher Scientific, Germany) at 60°C for 48 hours. Sample weight was recorded before and after drying and used to calculate the total dry matter. Following this, 0.5 g of the dried crushed insect biomass was transferred to a 2 mL plastic vial, and 1 mL of chloroform:methanol mixture (2:1, v/v) was added. The samples were kept in continuous agitation (Heidolph Multi Reax 115V, Heidolph Scientific Products GmbH, Schwabach, Germany) at 2,000 rpm for 24 h at room temperature and in darkness, and centrifuged at 20,800 x *g* (Eppendorf centrifuge 5817R, Eppendorf SE, Hamburg, Germany) for 3 min. After the centrifugation, the supernatant was collected and used for measuring the total lipid content and the soluble sugar. The precipitate obtained after the centrifugation was also collected and mixed with 1 mL of 0.1 M NaOH under constant agitation (Heidolph Multi Reax 115V, Heidolph Scientific Products GmbH, Schwabach, Germany) at 2,000 rpm for 2 hours at 80°C. Protein and glycogen contents were then measured on the resolubilised pellet.

To determine the lipid content, the phospho-vanillin test (Williams *et al*., 2011) was applied: 50 µL of the first supernatant (opportunely diluted with 98% ethanol to stay within the linear part of the calibration line) was heated at 100°C for 10 min to allow solvent evaporation, mixed with 40 µL of 98% sulphuric acid and heated again at 100°C for 10 min. After cooling down to room temperature, 960 µL of phospho-vanillin reagent (1.2 mg vanillin dissolved in 0.2 ml hot water (90°C) and 0.8 ml 85% phosphoric acid) was added and the mixture was incubated in dark at room temperature for 45 min. The absorbance was measured at 525 nm, and the lipid concentration was determined via a canola oil calibration curve.

Proteins were quantified using the Lowry method with modifications (Cuff *et al*., 2021): 250 µL of insect extract was diluted in 0.1 M NaOH, mixed with 225 µL of Lowry reagent A (2 g of potassium sodium tartrate and 100 g of sodium carbonate dissolved in 500 mL 1 M NaOH) and incubated at 50°C for 10 min. The mixture was allowed to cool down at room temperature and 25 µL of Lowry reagent B (2 g potassium sodium tartrate and 1 g of copper sulphate pentahydrate in 100 mL NaOH 1M) were added. After further incubation for 10 min at room temperature in the dark, 750 µL of Lowry reagent C (1 N Folin-Ciocalteau reagent) was added and the mixture was heated to 50°C for 10 min. The absorbance was measured at 650 nm and the protein content was determined via comparison to a calibration line based on defined concentrations of bovine serum albumin.

To determine carbohydrate content (soluble sugar and glycogen), 100 µL of diluted insect extract was mixed with 950 µL of anthrone reagent (1 mg of anthrone powder dissolved in 1 mL of 95% sulphuric acid) and incubated for 10 min at 92°C (Cuff *et al*., 2021; Skillman *et al*., 2018). The absorbance was recorded at 620 nm after the mixture reached room temperature. Corn starch was used as standard.

Chitin content was determined by mixing 10 mg of cricket sample with 0.5 mL of sulfuric acid 72% kept under constant agitation at room temperature for 90 min (Zamani *et al*., 2008). Obtained mixtures were then diluted with 8.2 mL of distilled water and incubated 1 hour at 121°C. Following the hydrolysis, 0.5 mL of each mixture was further mixed with 0.5 mL of 1M sodium nitrite solution (sample) or 0.5 mL of distilled water (blank). Solutions of ammonium sulfamate (12%, 0.5 mL), MBTH (0.5%, 0.5 mL) and ferric chloride (0.5%, 0.5 mL) were then added and the absorbance was measured at 650 nm. Commercial chitin from shrimps was used as standard.

#### Amino acid profile

The amino acid profile of the insects was determined following the methods of Psarianos *et al*. (2023) with modifications. In brief, 0.1 g of crushed dry (60°C, 48h) sample (in duplicate) was hydrolysed with 5 mL of 6N HCl for 24 hours at 110°C. The hydrolysed samples were cooled down, transferred into a 15 mL centrifuge tube and centrifuged (Eppendorf centrifuge 5810R, Eppendorf SE, Hamburg, Germany) at 3,200 x *g* for 10 min. The obtained solution was then filtered through a 0.22 µm PTFE syringe filter and 200 µL of hydrolysed sample were transferred to a 1.5 mL vial, dried in a vacuum centrifuge (Concentrator Plus, Eppendorf SE, Hamburg, Germany) at 95°C for 1 hour, and diluted in a dilution buffer (membraPure GmbH, Hennungsdorf, Germany). The diluted sample (0.02 mL) was then injected into an ARACUS Classic amino acid analyser (membraPure GmbH, Hennigsdorf, Germany), equipped with an auto sampler for automatic sample injection, a high-resolution sodium exchange chromatographic column (125 x 4 mm, 7 µm) and a reactor for post-column derivatisation with ninhydrin. Sample absorbance was recorded at 440 nm and 570 nm and the area under each chromatographic peak was integrated for amino acid quantification. Retention time was considered for amino acid identification.

#### Mineral profiling

The mineral profiles of diets and insects were determined via inductively coupled plasma mass spectrometry (ICP-MS) using dedicated protocols for each matrix. Diet samples were acid-digested using a MARS 6 microwave digestion system (CEM GmbH, Kamp-Lintfort, Germany) in accordance with Keil *et al*. (2020) and subsequently used for iron quantification applying the ICP-MS protocol described by Maares *et al*. (2025) on a PerkinElmer Nexion 1000 instrument (PerkinElmer Inc., Rodgau, Germany). Cricket samples (100 mg) were subjected to acidic microwave digestion (Mars 6, CEM, Kamp-Lintfort, Germany) using 65% HNO_3_ (Suprapur, Merck, Billerica, MA, USA) and 30% H_2_O_2_ (Merck, Billerica, MA, USA). Digestion was carried out with the following parameters: 600 W, 15 min ramp, 20 min at 200°C, yielding a clear solution. To validate digestion and metal quantification, certified reference materials (fish muscle (ERM-BB 422), single cell (BCR-274) and pig kidney (ERM-BB 186)) were subjected to the same protocol. The respective recoveries are listed in Supplementary Table S1. Metal content in insects was quantified using ICP-MS/MS (8800 ICP-tandem mass spectrometry (ICP-QQQ-MS, Agilent Technologies, Waldbronn, Germany). Experimental conditions for ICP-MS/MS measurements are listed in Supplementary Table S2.

#### Oxidative status

For the determination of the oxidative status of the insect biomass, Thiobarbituric acid (TBA) reactive species (TBARs) were quantified according to the method published in Ojha *et al*. (2024).

Specifically, approximately 2.5 g of cricket material was mixed with 10 mL of 5% (w/v) TCA solution and homogenised with an ultraturrax (IKA T25, IKA-Werke GmbH & Co. KG, Staufen, Germany) for 1 min. Samples were then centrifuged (Eppendorf centrifuge 5810R, Eppendorf SE, Hamburg, Germany) at 3,200 x *g* for 10 minutes and filtered through a Whatman No.4 filter paper. Three screw-top glass test tubes, each containing 3 mL of TBA 0.03 M solution were prepared, and 3 mL of the filtrated sample were added to each tube. Following a 1 h incubation at 100°C, the tubes were cooled to room temperature and then centrifuged at 3200 x *g* for 5 min. The absorbance was spectroscopically read at 532 nm with a SPECORD 210 Plus Double-beam spectrometer (Analytic Jena GmbH & Co., Germany) and TBARs were quantified by using a calibration line prepared with a TMP (1,1,3,3-tetramethoxypropane) 25 µM solution. TBARs level in each sample was expressed as µg malondialdehyde (MDA) equivalents/kg sample, calculated by applying the following formula (Schmedes and Holmer, 1989):

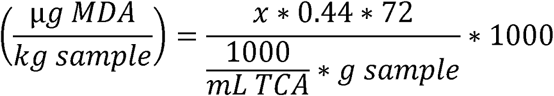

Where: *x* represents the MDA concentration(µM) in the sample expressed as TMP equivalents, 0.44 is the mass conversion factor between TMP and MDA (1 g of TMP is equivalent to 0.44 g of MDA), 72 (g/mol) is the molar mass of MDA, *mL TCA* is the volume of TCA used for sample hydrolysis, *g sample* accounts for the exact amount of sample used for the determination and the 1,000 factors are used for converting the volume of TCA from mL to L and the sample weight from g to kg.

### 2.3 Microbiome analysis

#### Microbial DNA extraction

Bacterial DNA was extracted from the final crickets as well as from the initial insect biomass (20 days old crickets). Before DNA extraction, samples were washed with 70% ethanol, rinsed twice with deionised water, placed in a 2 mL microcentrifuge tube and homogenised using a Qiagen TissueLyser II QIAGEN (Hilden, Germany) for 10 minutes at 25 Hz frequency. Total bacterial genomic DNA was extracted from the homogenised samples using Qiagen DNeasy PowerSoil Pro kits (Qiagen, Hilden, Germany) following the manufacturer’s protocol. In brief, approximately 250 mg of crushed sample were mixed with 800 µL of solution CD1 and agitated constantly for 10 min. The tubes were then centrifuged (15,000 rpm, 1 min), and 500 µL of supernatant was transferred into a new 2 mL microcentrifuge tube. 200 µL of solution CD2 was added, and the tubes were centrifuged again (15,000 rpm, 1 min). 700 µL of supernatant were then transferred to a new microcentrifuge tube containing 600 µL of solution CD3. 650 µL of the lysate was then loaded into a MB spin column, and the impurities were precipitated through a double centrifugation step (15,000 rpm, 1 min). After the second washing step, the MB spin column containing the extracted DNA was placed into a new collection tube, and 500 µL of EA solution was added to remove proteins and other non-aqueous contaminants. Clean DNA bound within the MB spin column was then purified once again with 500 µL of C5 solution and eluted in a vial with 100 µL of solution C6. Concentration (ng/mL) and purity (ratios A260/A280 and A260/A230) of the extracted DNA were determined with an Invitrogen Qubit 4.0 fluorometer (Thermo Fisher Scientific, USA) and a NanoDrop 2000 Spectrophotometer (Thermo Fisher Scientific, Waltham, MA, USA) respectively. The extracted DNA was stored at −20°C until further processing.

#### PCR amplification and 16S rRNA sequencing

The bacterial 16S rRNA gene amplification targeted the 16S rRNA V3-V4 region using universal primer set 515F (5’-TCGTCGGCAGCGTCAGATGTGTATAAGAGACAGGTGYCAGCMGCCGCGGTA-3’) and 806R (5’-GTCTCGTGGGCTCGGAGATGTGTATAAGAGACAGGGACTACNVGGGTWTCTAAT-3’). Polymerase chain reaction (PCR) consisted in an initial denaturation at 98°C for 1 min, followed by 30 cycles of denaturation at 98°C for 10 s, annealing at 50°C for 30 s and elongation at 72°C for 30 s. A final elongation step at 72°C for 5 min was also carried out. PCR products were purified to remove primer dimers and non-specific products before indexing and library preparation. Sequencing libraries were prepared as described in Antunes *et al*. (2025), where all pipetting and library preparation steps were automated using the Beckman Coulter Biomek i7 Automated Liquid Handler (Beckman Coulter Life Sciences, Indianapolis, IN, USA). Libraries were subjected to paired-end sequencing on the Illumina MiSeq platform using 2 × 250 bp as described in Antunes *et al*. (2025). Raw sequencing reads were processed using the QIIME2 platform.

Sequences were trimmed using cutadapt in python, then analysed using the R packages *dada2* (Callahan *et al*., 2016) and *phyloseq* (McMurdie and Holmes, 2012), following the tutorials provided by the packages authors (https://benjjneb.github.io/dada2/tutorial.html; https://rpubs.com/telatin/day3-tutorial) Sequence processing included demultiplexing, quality filtering, removal of chimeric sequences and ambiguous reads, and denoising to generate Amplicon Sequence Variants (ASVs).

Taxonomic assignment of representative sequences was performed against the Greengenes database to generate taxonomic abundance tables at phylum, class, and genus levels. Absolute abundance was then converted into relative abundance and data of different pseudoreplicates belonging to the same sample were combined. Microbial alpha diversity was evaluated using the Shannon Diversity Index and Simpson Diversity Index to assess microbial richness and evenness within the samples. Beta diversity analysis was performed using the Bray–Curtis dissimilarity matrix as well as the Jaccard dissimilarity matrix, aiming to evaluate the differences in microbial community composition among different treatment groups.

### 2.4 Statistical analyses

All statistical analyses were performed in R (version 4.5.1). Chemical as well as rearing performance data were checked for normality and homogeneity of variance (homoscedasticity) using the Shapiro-Wilk and the Levene’s tests, respectively. When the assumptions were fulfilled, a one-way analysis of variance (ANOVA) at probability level of 0.05 was applied. When the normality assumption was not fulfilled, data were normalized via Box-Cox transformation. One-way ANOVAs followed by Tukey’s HSD tests (p-value = 0.05) were then carried out. Insect survival was analysed via a general linear model (GLM), with a binomial distribution (link “logit”). Average cricket daily weight was analysed via GLM, supposing a Gamma distribution (link “log”) as suggested by Pisa *et al*. (2025). A GLMM (generalized linear mixed model) with Gamma distribution (link “log”) and data nested within the same box was instead used for studying the overall growth curves (Oddon *et al*., 2024; Pisa *et al*., 2022). Differences on the overall amino acid profiles were evaluated through a Principal Component Analysis (PCA) followed by permutational multivariate analysis of variance (PERMANOVA) with 999 permutations. Hotteling pairwise T2 post-hoc tests at probability level of 0.05 was applied for confirming the differences between treatment groups.

Microbial alpha diversity metrics were compared between treatments using linear models (LM). Shapiro-Wilk test and Levene’s test were computed for normality and homoscedasticity tests. Non-parametric Kruskal-Wallis test was implemented when model assumptions were not respected. Beta Diversity matrixes were visualized through Principal Coordinate Analysis (PCoA), and analysed using PERMANOVA with 9,999 permutations. Potential variations in relative abundance of single taxa were visually evaluated through heat maps built at class and genus levels, while Linear Discriminant Effect Size (LEfSe) analysis with a LDA cut-off of 2 and 1 and a significant level of 0.05 were computed for identifying potential microbial biomarkers.

## 3 Results

### 3.1. Rearing and bioconversion performances

Over 21 days of-feeding trials (Supplementary Figure S1), dietary iron supplementation did not significantly affect *A. domesticus* growth (GLMM with gamma distribution, χ = 4.584, p = 0.4687), with growth curves showing a similar trend, regardless by the treatment (Figure 1a). The fastest growth was always observed between day 3 and day 17, and the curves tended to flatten out in the following days, suggesting that the metamorphosis phase was reached. However, although no statistically significant differences were detected, crickets reared on diets Fe_5mM, Fe_10mM, Fe_25mM and Fe_100mM reached higher weights on day 19, while crickets from the treatment Fe_1mM stopped growing at day 17 and crickets from the CTR treatments showed slightly slower and prolonged growth. Crickets’ survival did, however, differ significantly among treatments (GLM with binomial distribution, p = 0.0112) with the Fe_5mM showing the highest survival (96.42 ± 1.47 %) and treatments CTR (88.45 ± 0.95 %), Fe_25mM (88.00 ± 2.10 %) and Fe_100mM (86.88 ± 2.44 %) the lowest (Figure 1b). No dose dependent effect of iron supplementation was observed.

**Figure 1.**
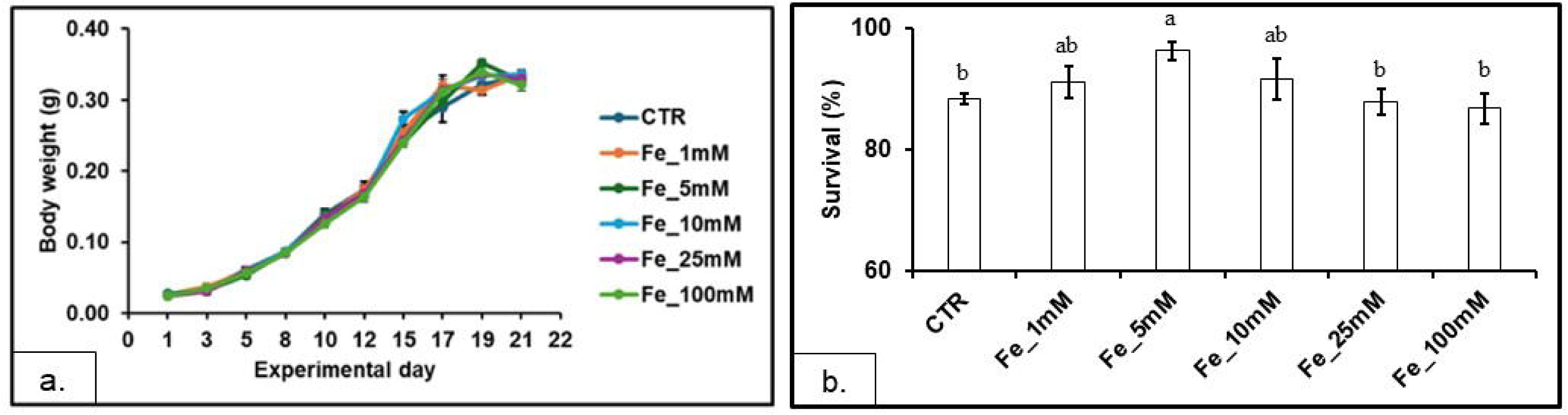
Growth curves (a.) and survival (b.) of crickets reared on diets supplemented with different level of iron. Both measurements were recorded starting from 20-days-old *A. domesticus* nymphs over 21 days of rearing. Survival was monitored throughout the experimental period, during which crickets were fed experimental diets spiked with different concentrations of FeSO_4_. Values are presented as mean ± SE (n = 5). For the bar plot, different letters on top of the bars indicate statistically significant differences (one-way ANOVA followed by Tukey’s HSD post hoc test; P < 0.05).

In contrast to growth and survival, feed conversion ratio (FCR; F_(5,24)_ = 16.22; p < 0.001) and total feed intake (F_(5,24)_ = 14.58; p < 0.001) showed a dose-dependent response to dietary iron supplementation (Table 2). In both cases, increasing the Fe in the diet led to a progressive reduction in feed intake and FCR, with the CTR group displaying values statistically similar to the Fe_5mM, Fe_10mM and Fe_25mM treatments. The worst performances (higher FCR and lower feed intake) were recorded in the group Fe_1mM, while the group Fe_100mM stood out as the best performing treatment. No differences were detected in the Average Daily Gain (ADG, F_(5,24)_ = 0.623; p = 0.684), which remained stable across the treatment groups, ranging between 14.17 ± 0.39 g (Fe_100mM) and 14.78 ± 0.34 g (Fe_10mM).

**Table 2.**
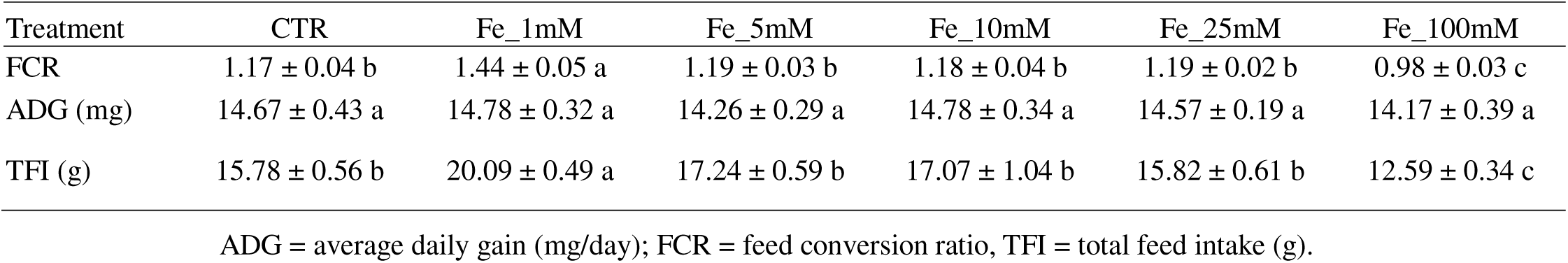
Growth and feed conversion performances of *A. domesticus* nymphs fed with experimental diets spiked with increasing iron concentrations. Data are reported as mean ± SE (n = 5). Different letters within the same column indicate statistically significant differences (one-way ANOVA followed by Tukey’s HSD post hoc test; P < 0.05).

| Treatment | CTR | Fe_1mM | Fe_5mM | Fe_10mM | Fe_25mM | Fe_100mM |
|---|---|---|---|---|---|---|
| FCR | 1.17 ± 0.04 b | 1.44 ± 0.05 a | 1.19 ± 0.03 b | 1.18 ± 0.04 b | 1.19 ± 0.02 b | 0.98 ± 0.03 c |
| ADG (mg) | 14.67 ± 0.43 a | 14.78 ± 0.32 a | 14.26 ± 0.29 a | 14.78 ± 0.34 a | 14.57 ± 0.19 a | 14.17 ± 0.39 a |
| TFI (g) | 15.78 ± 0.56 b | 20.09 ± 0.49 a | 17.24 ± 0.59 b | 17.07 ± 1.04 b | 15.82 ± 0.61 b | 12.59 ± 0.34 c |

ADG = average daily gain (mg/day); FCR = feed conversion ratio, TFI = total feed intake (g).

### 3.2 Chemical analysis

No significant differences were observed on the proximate composition of crickets, with values of proteins (F_(5,24)_ = 0.746; p = 0.597), lipids (F_(5,24)_ = 0.601; p = 0.700), soluble carbohydrates (F_(5,24)_ = 0.952; p = 0.466) and glycogen (F_(5,24)_ = 1.712; p < 0.170) being always statistically comparable between groups (Table 3). Significant differences were, however, detected for dry matter (F_(5,24)_ = 3.058; p < 0.0283) and chitin (F_(5,24)_ = 9.306; p < 0.0001), with the last increasing when higher iron levels were included in the diet. Indeed, although chitin levels ranged between 10.62 ± 0.65 % dry matter (Fe_1mM) and 19.97 ± 2.21 % dry matter (Fe_100mM), they were lower in the CTR and Fe_1mM groups, slightly increasing in the Fe_10mM (15.38 ± 0.53 % dry matter) and Fe_25mM (15.81 ± 0.45 % dry matter) and reaching the maximum level in the Fe_100mM crickets. No significative differences were, however, observed between the Fe_100mM and the Fe_5mM groups (Figure 2a) suggesting absence of direct correlation between Fe dietary inclusion and chitin content. On the other hand, a clear dose-dependent effect was observed for thiobarbituric acid reactive substances (TBARs, Figure 2b), which did not differ at the iron concentration treatments up to Fe_25mM (120.72 – 136.55 µg MDA/kg fresh matter), but were significantly higher in the Fe_100mM treatment (199.94 ± 14.51 µg MDA/kg fresh matter; F_(5,24)_ = 7.857; p = 0.0002).

**Figure 2.**
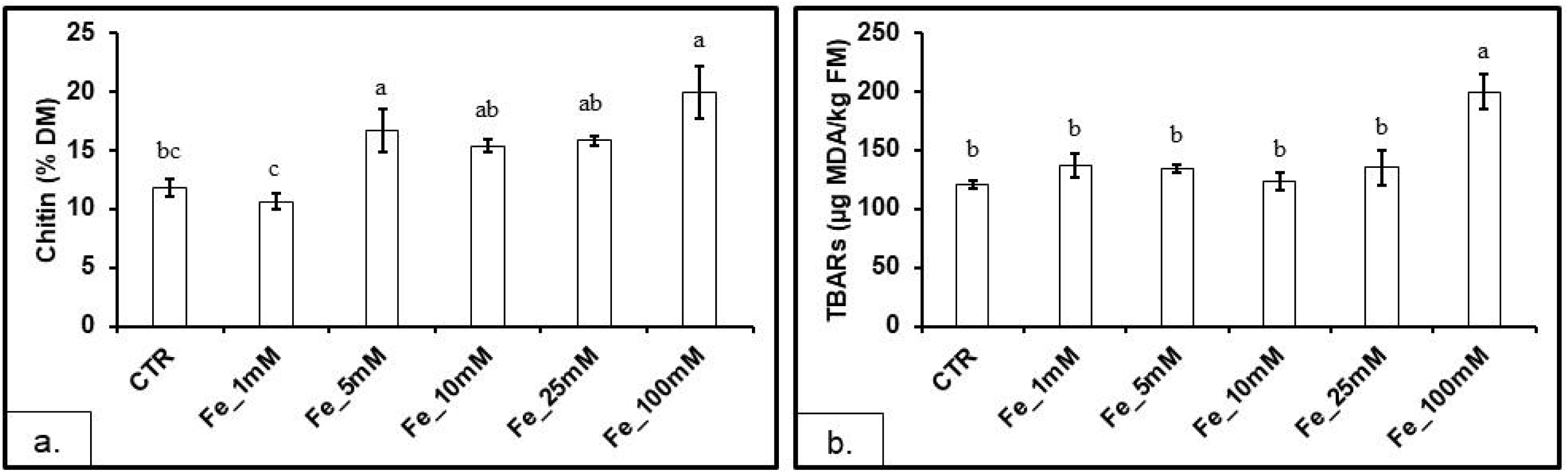
Chitin level (a.) and TBARs (b.) of *A. domesticus* nymphs reared on diets supplemented with different levels of iron. Values are presented as mean ± SE (n = 5). Different letters within the same plot indicate statistically significant differences (one-way ANOVA followed by Tukey’s HSD post hoc test; P < 0.05).

**Table 3.** Proximate analysis of *A. domesticus* nymphs fed with experimental diets spiked with increasing iron concentrations. Data are reported as mean ± SE (n = 5). Different letters within the same row indicate statistically significant differences (one-way ANOVA followed by Tukey’s HSD post hoc test; P < 0.05).

| Treatment | CTR | Fe_1mM | Fe_5mM | Fe_10mM | Fe_25mM | Fe_100mM |
|---|---|---|---|---|---|---|
| Dry Matter (% FM) | 26.41 ± 0.26 a | 24.81 ± 0.26 b | 26.12 ± 0.50 ab | 25.54 ± 0.21 ab | 25.47 ± 0.51 ab | 26.29 ± 0.21 ab |
| Protein (% DM) | 50.37 ± 0.51 a | 51.25 ± 1.07 a | 49.55 ± 1.41 a | 50.70 ± 0.26 a | 50.08 ± 0.96 a | 49.00 ± 1.08 a |
| Lipids (% DM) | 34.11 ± 1.09 a | 34.20 ± 1.92 a | 36.50 ± 1.45 a | 33.85 ± 1.17 a | 33.37 ± 1.52 a | 33.48 ± 1.59 a |
| Glycogen (% DM) | 0.60 ± 0.04 a | 0.58 ± 0.05 a | 0.59 ± 0.03 a | 0.66 ± 0.02 a | 0.55 ± 0.02 a | 0.66 ± 0.03 a |
| Soluble carbohydrates (% DM) | 0.53 ± 0.02 a | 0.54 ± 0.02 a | 0.58 ± 0.03 a | 0.54 ± 0.02 a | 0.53 ± 0.02 a | 0.52 ± 0.02 a |

Significant differences were observed in the amino acid profile of the crickets (Supplementary Table S3). Increased iron level in diet led to a progressive linear increase of aspartic acid (F_(5,24)_ = 13.45; p < 0.0001), threonine (F_(5,24)_ = 19.37; p < 0.0001), serine (F_(5,24)_ = 15.28; p < 0.0001) and glutamic acid (F_(5,24)_ = 14.08; p < 0.0001) in crickets. Other amino acids, including alanine (F_(5,24)_ = 17.00; p < 0.0001), valine (F_(5,24)_ = 20.32; p < 0.0001), methionine (F_(5,24)_ = 21.78; p < 0.0001), leucine (F_(5,24)_ = 15.86; p < 0.0001), isoleucine (F_(5,24)_ = 17.44; p < 0.0001), tyrosine (F_(5,24)_ = 23.46; p < 0.0001), histidine (F_(5,24)_ = 15.72; p < 0.0001), phenylalanine (F_(5,24)_ = 7.762; p = 0.0002), lysine (F_(5,24)_ = 19.16; p < 0.0001) and arginine (F_(5,24)_ = 18.14; p < 0.0001), displayed a singular behaviour, with high levels observed in the CTR crickets, followed by a reduction in the Fe_1mM treatment, a progressive linear increase between the Fe_1mM and the Fe_25mM groups and a slight reduction in the Fe_100mM crickets. No differences were observed for proline (F_(5,24)_ = 2.472; p = 0.0609), while glycine (F_(5,24)_ = 11.80; p < 0.0001) peaked in the Fe_100mM treatment, showing significantly lower concentration in the Fe_25mM group. The principal component analysis biplot built on the amino acid profiles (Figure 3) confirmed these differences (PERMANOVA: F = 14.727, p = 0.001; beta dispersion: p = 0.941) suggesting that increased amount of iron in diet led to crickets enriched of serine, aspartic acid, threonine, glutamic acid, alanine, proline and glycine, while tyrosine, valine, leucine, isoleucine, arginine, lysine, methionine, histidine and phenylalanine characterized the Fe_25mM crickets, which grouped separately from all the others.

**Figure 3.**
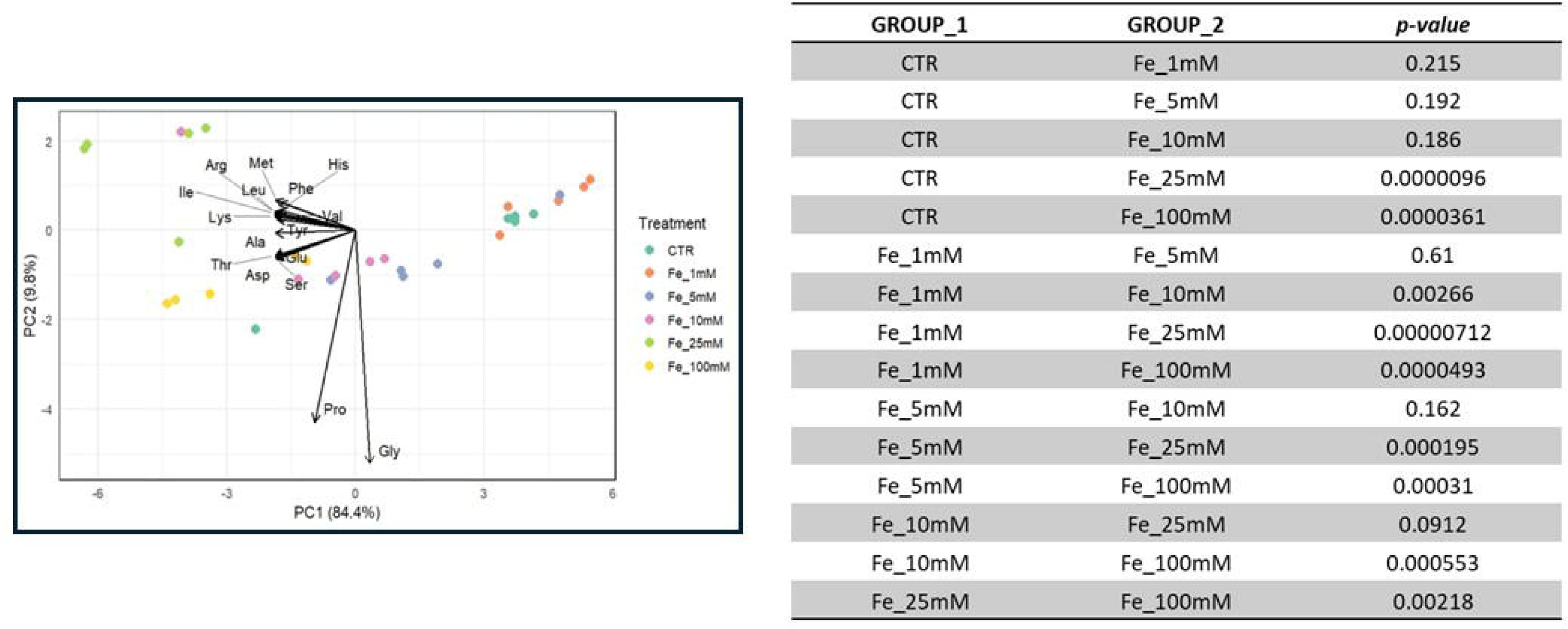
PCA biplot built on the amino acid profiles of *A. domesticus* nymphs reared on diets supplemented with different levels of iron. Values in the table represent the results of the Hotelling pairwise T2 post hoc test run after PERMANOVA on the same amino acid data.

Dietary iron supplementation led to a clear dose-dependent accumulation of iron in crickets, while leaving other minerals unaffected (Table 4). Most of the analysed metals did not differ significantly among groups (p > 0.05). However, the iron content of the crickets increased up to five times (F_(5,24)_ = 22.02; p < 0.0001) with levels of 21.38 ± 1.08 mg/kg dry matter in the CTR group (BAF_Fe_ ∼ 0.06) compared to 118.34 ± 55.60 mg/kg dry matter in the Fe_100 mM group (BAF_Fe_ ∼ 0.02).

**Table 4.** Macro- and trace-mineral composition of *A. domesticus* nymphs reared on diets supplemented with different levels of iron. Results are presented as mean ± SE (n = 5). Different letters within the same row indicate statistically significant differences (one-way ANOVA followed by Tukey’s HSD post hoc test; P < 0.05).

| Treatment | CTR | Fe_1mM | Fe_5mM | Fe_10mM | Fe_25mM | Fe_100mM |
|---|---|---|---|---|---|---|
| Fe (mg/kg DM) | 21.38 ± 1.08 c | 23.30 ± 0.96 c | 28.62 ± 2.84 bc | 34.21 ± 2.18 b | 41.09 ± 6.53 b | 118.34 ± 55.60 a |
| Zn (mg/kg DM) | 63.62 ± 3.08 a | 64.46 ± 6.62 a | 53.90 ± 3.50 a | 67.77 ± 6.62 a | 55.24 ± 3.78 a | 50.23 ± 1.38 a |
| Cu (mg/kg DM) | 8.05 ± 0.60 a | 5.96 ± 0.31 a | 6.84 ± 0.81 a | 6.57 ± 0.93 a | 6.26 ± 0.35 a | 6.52 ± 0.53 a |
| Mn (mg/kg DM) | 7.89 ± 0.35 a | 7.13 ± 0.56 a | 7.04 ± 0.41 a | 6.65 ± 0.60 a | 5.52 ± 0.49 ab | 4.00 ± 0.99 b |
| Se (µg/kg DM) | 163.72 ± 64.19 a | 123.01 ± 20.37 a | 138.17 ± 36.00 a | 131.87 ± 28.35 a | 103.63 ± 7.37 a | 103.72 ± 3.60 a |
| Mo (µg/kg DM) | 159.76 ± 4.94 a | 156.19 ± 8.80 a | 158.25 ± 3.43 a | 170.40 ± 9.28 a | 170.01 ± 21.87 a | 149.06 ± 4.39 a |
| Co (µg/kg DM) | 8.40 ± 1.28 a | 8.51 ± 0.82 a | 12.09 ± 2.59 a | 12.82 ± 1.94 a | 12.24 ± 4.49 a | 13.92 ± 3.96 a |
| Ca (g/kg DM) | 0.32 ± 0.01 a | 0.32 ± 0.03 a | 0.30 ± 0.02 a | 0.36 ± 0.04 a | 0.31 ± 0.04 a | 0.31 ± 0.03 a |
| P (g/kg DM) | 2.09 ± 0.03 a | 1.98 ± 0.06 a | 1.92 ± 0.10 a | 2.23 ± 0.17 a | 2.02 ± 0.12 a | 1.89 ± 0.03 a |
| K (g/kg DM) | 2.76 ± 0.07 a | 2.67 ± 0.07 a | 2.62 ± 0.10 a | 3.01 ± 0.19 a | 2.75 ± 0.10 a | 2.72 ± 0.07 a |
| Na (g/kg DM) | 1.07 ± 0.01 a | 1.01 ± 0.03 a | 0.99 ± 0.04 a | 1.16 ± 0.08 a | 1.01 ± 0.05 a | 1.05 ± 0.02 a |
| S (g/kg DM) | 1.39 ± 0.02 a | 1.31 ± 0.03 a | 1.30 ± 0.07 a | 1.45 ± 0.07 a | 1.34 ± 0.06 a | 1.26 ± 0.03 a |
| Mg (g/kg DM) | 0.21 ± 0.00 a | 0.20 ± 0.01 a | 0.20 ± 0.01 a | 0.24 ± 0.02 a | 0.20 ± 0.01 a | 0.19 ± 0.00 a |
| Cd (µg/kg DM) | 13.62 ± 2.43 a | 12.45 ± 2.30 a | 13.04 ± 1.25 a | 14.64 ± 2.39 a | 10.19 ± 1.26 a | 8.24 ± 0.83 a |
| Pb (µg/kg DM) | 28.96 ± 11.91 a | 23.18 ± 2.98 a | 25.46 ± 2.59 a | 29.29 ± 6.89 a | 17.47 ± 4.51 a | 25.86 ± 4.84 a |
| As (µg/kg DM) | 3.18 ± 0.22 a | 3.15 ± 0.38 a | 3.55 ± 0.31 a | 3.54 ± 0.31 a | 3.22 ± 0.37 a | 4.07 ± 0.48 a |

Interestingly, manganese showed a significant decrease from the CTR (7.89 ± 0.35 mg/kg) to the Fe_100 mM (4.00 ± 0.99 mg/kg) crickets, with a very strong dose-dependent relationship (F_(5,24)_ = 5.338; p < 0.0020). No differences were observed in the toxic elements, with arsenic (F_(5,24)_ = 0.995; p = 0.441) and lead (F_(5,24)_ = 0.447; p = 0.244) staying stable regardless of the dietary treatment. Notably, cadmium levels showed a non-significant trend towards dose-dependent decline, with the Fe_25mM and Fe_100mM groups displaying lower concentrations than the remaining treatments; however, these differences were not statistically significant (F_(5,24)_ = 1.636; p = 0.189).

### 3.3 Microbiome analysis

The cricket microbiome showed a consistent high alpha diversity across treatments, suggesting high number of different microbial taxa in every sample, regardless of the treatment. The Shannon index ranged between 4 and 4.8, with the lowest values recorded in the starter crickets and higher values computed on the harvested insects (Figure 4a), indicating an increase in microbiome diversity with age. Simpson diversity, which considers the evenness of the microbial taxa, was characterized by even higher stability, ranging consistently between 0.96 and 0.99 (Figure 4b). There were no significant differences among treatments in either diversity index (Kruskal-Wallis-Test: Shannon diversity: H_(6)_ = 8.7273, p = 0,1895; Simpson diversity: H_(6)_ = 7.5009, p = 0.277).

**Figure 4.**
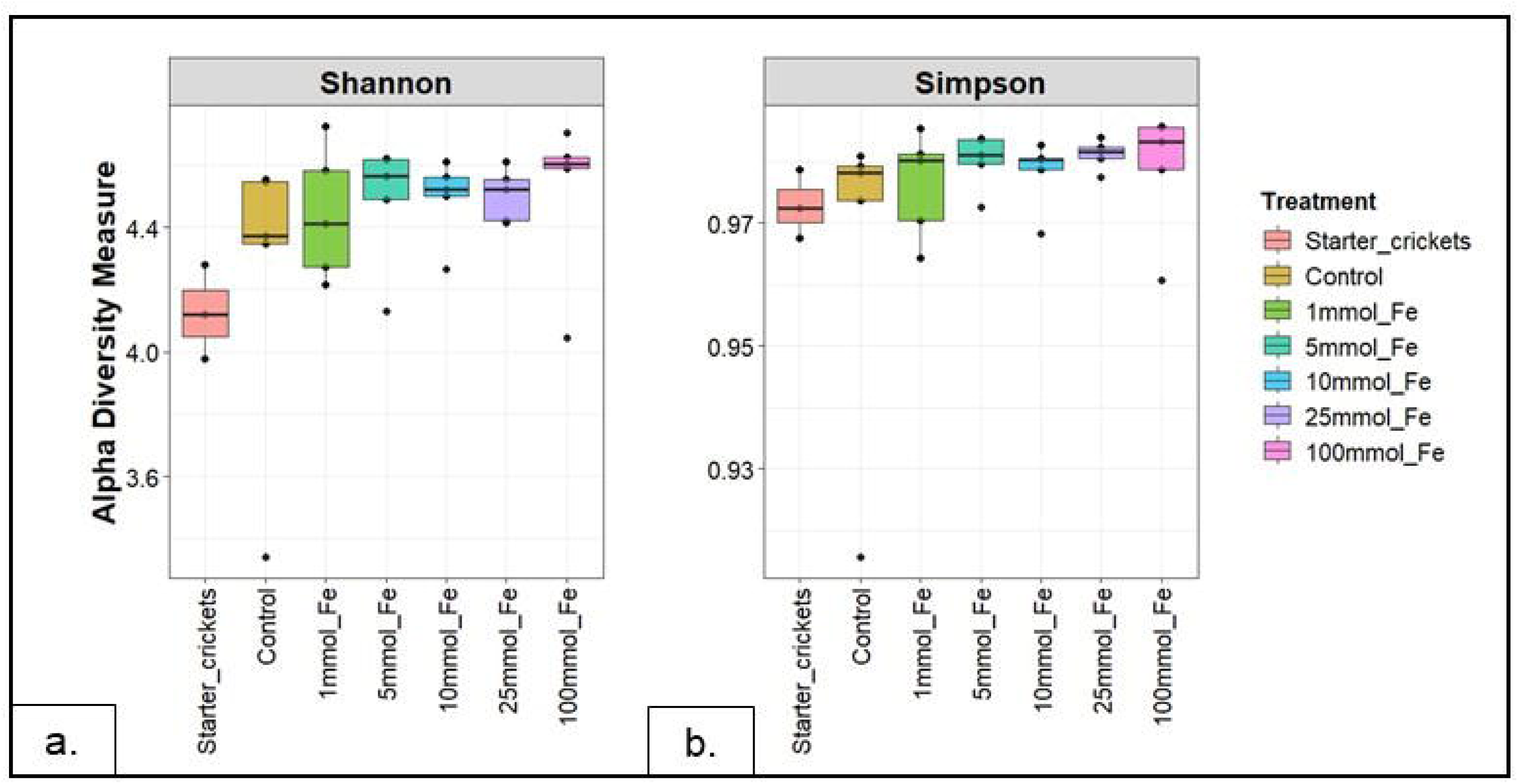
Alpha diversity (a.: Shannon diversity index; b.: Simpson diversity index) of *A. domesticus* nymphs reared on diets supplemented with different levels of iron and starter crickets.

The beta diversity results built on the Bray-Curtis dissimilarity matrix supported the alpha diversity findings, indicating that different levels of iron in diet did not lead to significant differences in terms of microbial community composition (PERMANOVA: F_(5)_ = 0.7935, p = 0.94). However, we did observe a significant microbial community shift between juvenile and adult crickets (F_(6)_ = 1.3959, p = 0.007). The PCoA run on the Bray-Curtis dissimilarity matrix (Figure 5a) clearly showed these observations, with samples from different dietary treatments clustering together, but separately from the starter crickets. Similar results were also obtained when the beta diversity analysis were carried out using the Jaccard dissimilarities matrix (PERMANOVA without starter crickets: F_(5)_ = 0.8671, p = 0.93; PERMANOVA with starter crickets: F_(6)_ = 1.3321, p = 0.002) (Figure 5b), indicating that the differences between starter crickets and harvested crickets and the similarities between crickets grown at different iron levels were not only due to the taxon relative abundance, but also to the presence/absence of the different microbial taxa. Results of the pairwise PERMANOVAs can be found in Supplementary Table S4.

**Figure 5.**
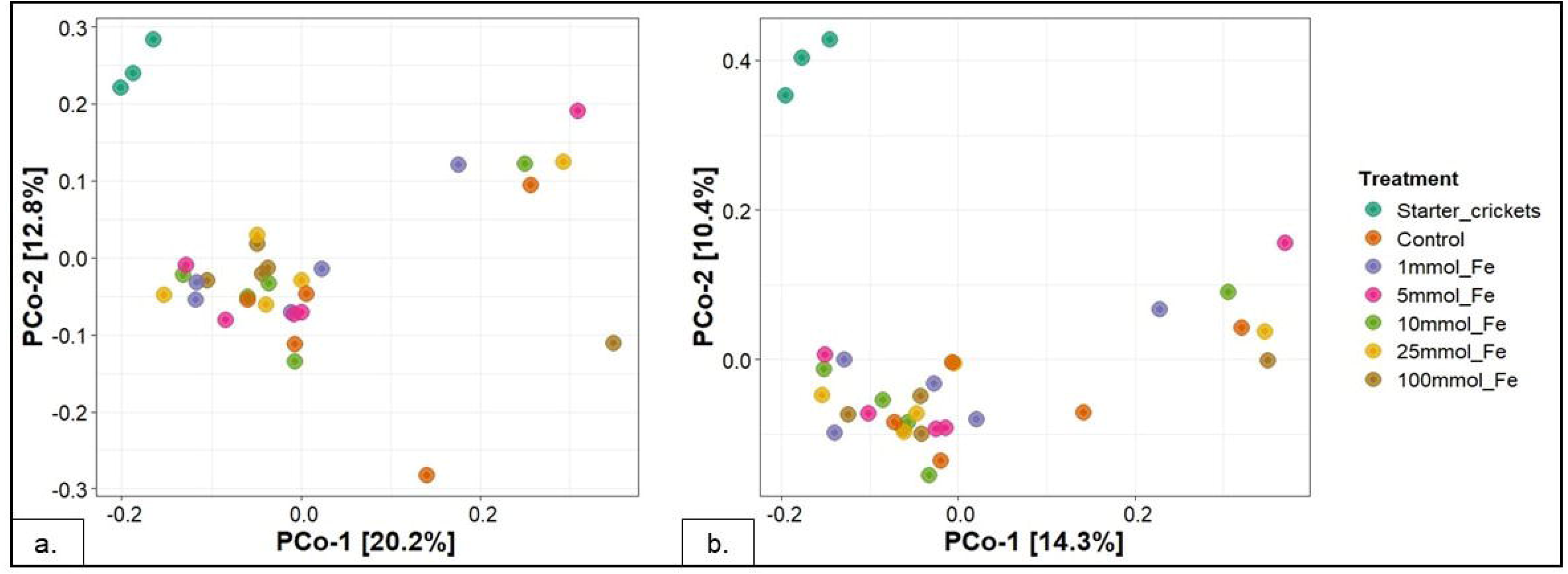
PCoA built on the beta diversity indexes (a.: computed by using the Bray-Curtis dissimilarity matrix; b.: computed by using the Jaccard dissimilarity matrix) of *A. domesticus* nymphs reared on diets supplemented with different levels of iron and starter crickets.

At the phylum level, the whole-body cricket microbiome was predominantly composed of Firmicutes, Bacteroidota and Proteobacteria (Figure 6a), which remained stable across treatments. Verrucomicrobiota, Desulfobacterota, Deferribacterota and Actinobacteriota also shaped the core microbiota of the crickets with a consistently high abundance across samples, although these phyla were present at lower abundancies than the Firmicutes, Bacteroidota and Proteobacteria. At the class level (Figure 6b), the most abundant taxa were Clostridia (phylum Firmicutes), Bacteroidia (phylum Bacteroidota), Gammaproteobacteria (phylum Proteobacteria) and Bacilli (phylum Firmicutes), which appeared stable among treatments. Lower abundance was detected for Acidimicrobiia (phylum Actinobacteriota), Verrucomicrobiae (phylum Verrucomicrobiota), Desulfovibrionia (phylum Desulfobacterota) and Coriobacteriia (phylum Actinobacteriota), which, although still part of the cricket core microbiome, showed higher variability.

**Figure 6.**
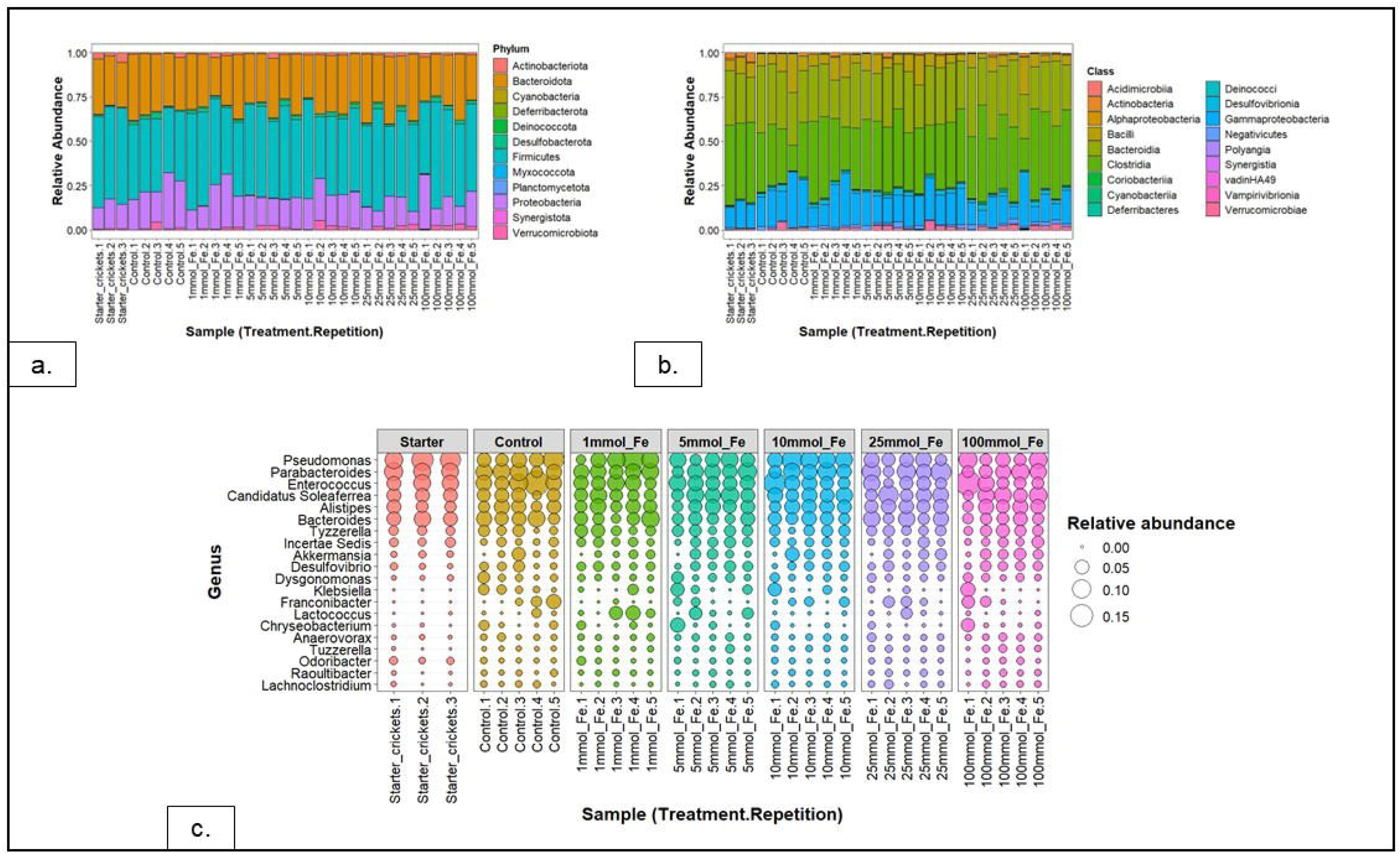
Relative abundance computed at (a.) phylum level, (b.) class level, (c.) genus level of the whole-body microbiome of *A. domesticus* nymphs reared on diets supplemented with different levels of iron and starter crickets.

The high consistency of the cricket microbiome among treatments is also confirmed at the genus level (Figure 6c). *Pseudomonas* spp. (phylum Proteobacteria), *Parabacteroides* spp. (phylum Bacteroidota), *Dysgonomonas* spp. (Bacteriodota) and *Enterococcus* spp. (phylum Firmicutes) appear to be the most abundant genera in all the samples. However, when comparing the starter crickets with the harvested ones, it appears that the first have lower relative abundance in Enterobacterales-associated genera such as *Klebsiella* spp. and *Franconibacter* spp., and higher abundance in Firmicutes such as Lachnospiraceae and *Enterococcus* spp., and Bacteroidota genera such as *Dysgonomonas* spp..

Consistently with these analyses, no microbial biomarkers were identified in any sample, even after applying the very strict LEfSE cut-off value of 1. However, differences in relative abundance of the different microbial taxa were noticed at both, class and genus levels, suggesting that, although the microbial composition was quite stable among treatments, provision of diets with different iron level can still affect the organization of the overall insect microbial community.

## 4. Discussion

This study aimed to investigate the impact of high dietary iron concentrations on the growth, survival and nutritional composition of *A. domesticus*. We evaluated how rearing under iron excess shapes the insects’ macronutrient composition, microbiome, and multielement profile. A central finding of this study is the remarkable stability of both the core nutritional parameters (total protein, carbohydrates, glycogen and lipids), the microbiome and the phenotypic growth performance of *A. domesticus* under dietary iron enrichment. Crucially, even under severe iron overload of ∼ 6,900 mg Fe/kg diet (Fe_100mM), there were no consistent reductions in either final body weight or overall survival. This resilience demonstrates exceptionally robust homeostatic control, with crickets avoiding the disruption of vital physiological functions under massive iron supply. This aligns with the geometric framework of nutrition, which posits that insects actively regulate nutrient intake to maintain a defined assimilation target and thereby stabilize internal body composition despite dietary imbalances (Harrison *et al*., 2014; Rho and Lee, 2022).

Survival in the Fe_100mM group was significantly lower than in the lower iron treatments (1–10 mM), but it did not differ from the control group and remained overall high (around 90%). This suggests that while higher iron doses induced a physiological response, they did not cause acute lethality. In fact, in general, the survival levels we observed exceed those typically reported in literature, where values below 80% are common for *A. domesticus* (Oonincx et al., 2015; Mahavidanage et al., 2023). A plausible explanation for the decline in survival of the Fe_100mM group relative to the lower supplementation levels (1–10 mM) might be a saturation of the insect’s iron storage and detoxification capacity. At extreme dietary doses, the physiological mechanisms responsible for iron sequestration—such as ferritin binding—are likely overwhelmed. This non-linear handling of high iron influx can lead to an accumulation of free, unsequestered iron in the gut, a process previously described in other insects, including *Aedes aegypti* and *H. illucens* larvae (First *et al*., 2026; Geiser *et al*., 2009; Geiser *et al*., 2006). This free iron can readily participate in a Fenton reaction, potentially inducing cellular oxidation (Cardoso-Jaime *et al*., 2022; Dunkov and Georgieva, 2006; Weber *et al*., 2020). This hypothesis is strongly supported by the elevated TBARS levels observed in the Fe_100mM crickets. Conversely, the peak in survival observed at the moderate Fe_5mM dose points toward an optimal physiological window for iron supplementation. Iron is an essential micronutrient required for numerous core metabolic pathways, and providing a well-balanced supply can directly enhance overall insect fitness, metabolic health, and baseline immune resilience. At this moderate level, the influx of iron likely matches the insect’s natural homeostatic capacity, ensuring that the element is efficiently utilized and sequestered without causing the gut accumulation or oxidative stress observed at higher doses. This optimal systemic balance may sustain robust physiological barriers and immunological functions, thereby contributing to the enhanced vitality and survival observed in this treatment group.

In general, the physiological effects of iron supplementation in mass-reared insects vary considerably across species. Iron supplementation of diets with ammonium citrate at a final concentration of 563 mg Fe/kg diet led to 4% lower survival in *T. molitor* larvae, although even higher concentrations (up to 6,637 mg Fe/kg diet) did not affect *H. illucens* larvae survival (First *et al*., 2024; First *et al*., 2026). Presence of cuticular lesions indicative of iron overload was however recorded in black soldier fly larvae kept at such a high iron concentration (First *et al*., 2024). Taken together, these findings suggest that the physiological responses to dietary nutrients in insects are not standardizable, being highly dependent on both the specific insect species and the underlying dietary regime (Bawa *et al*., 2020; Gorman, 2023; Wang *et al*., 2026).

Interestingly, in this study’s *A. domesticus* groups, the survival outcomes under dietary iron enrichment were not associated with any changes in the whole-cricket microbiome. Microbiome composition, along with alpha and beta diversities were in fact remarkably consistent across all treatments. This observation is reassuring from a safety perspective, as we saw neither a proliferation of pathogens nor any evidence of dysbiosis in treated crickets. Previous research on the impact of iron supplementation on insect microbiome composition has shown mixed results. In a recent study on the mosquito *Anopheles culifacensis,* Yadav *et al*. (2026) observed a selective shift in the gut microbiota following exposure to a 10 mM ferric chloride-enriched diet. In this case there was a clear taxonomic restructuring in response to iron treatment, with iron-tolerant bacterial strains progressively proliferating in the presence of increasing iron availability, while vulnerable taxa were suppressed. This is a phenomenon that has also been observed in vertebrates (Malesza *et al*., 2022; Parmanand *et al*., 2019). Our results are, however, consistent with findings from other Orthopterans, including *Acheta domestisticus*, where the microbiome continues to support nutrient uptake, immune function, and stress tolerance under conditions of iron supplementation up to 100 mmol Fe (Hance *et al*., 2025; Shamjana *et al*., 2024). The absence of remarkable microbial changes observed in our study combines with previous findings from Orthopterans suggests that the microbiome of Orthopterans is particularly resilient to iron supplementation.

The fact that high phenotypic tolerance in crickets was accompanied by a trend toward lower feed conversion ratios suggests that dietary iron might have enhanced metabolic efficiency by optimizing iron-dependent mitochondrial respiration (cytochromes) and activity of crucial enzymes like catalase, directly involved in iron detoxification (Gorman, 2023; Oliveira *et al*., 2017). However, despite this efficiency, total feed intake significantly declined at higher iron doses, indicating behavioural feed avoidance due to iron-induced appetite disruption or palatability shifts (Gekière, 2025; Mogren and Trumble, 2010). Similar effects have been observed in black soldier fly larvae reared on fresh aquaculture sludge (Rossi *et al*., 2023) and might be due to a compensatory mechanism whereby insects tend to consume less substrate (lower feed intake), maximizing its use (lower FCR). This behavioural restriction likely helped to limit systemic metal influx. Nevertheless, the iron load might still have altered the redox environment inside the crickets, as suggested by the sharp increase in TBARs in the Fe_100mM treatment. Indeed, at these excessive exposure levels, any labile iron that might accumulate in the midgut could trigger Fenton-type reactions, producing destructive hydroxyl radicals (Barbehenn *et al*., 2005; Barbehenn and Stannard, 2004; Walter-Nuno *et al*., 2018) and triggering a systemic lipid peroxidation cascade that extends beyond the gut into peripheral tissues. Specific structural barriers, such as the external cuticular matrices and the semipermeable peritrophic membrane of the midgut, might therefore act to restrict excessive metal accumulation thus preventing massive toxic influxes (Gekière, 2025). Specifically, as the structural chitin contained in the luminal gut possesses high iron-binding capacity (Anastopoulos *et al*., 2017; Lehnert *et al*., 2025), it might act as an iron storage pool with an analogous function to human mucins (Einhorn *et al*., 2024), minimizing the pool of free iron potentially involved in oxidative reactions. Such an idea is supported by the increase in chitin content observed at high iron levels (treatment Fe_10mM). In addition to chitin, the observed increase of tyrosine under high iron exposure further substantiates this coordinated cuticular defence mechanism. Tyrosine is the foundational amino acid required for sclerotization and melanisation in insects (Gorman and Arakane, 2010). These structural tanning processes not only harden the exoskeleton but also embed the chelated iron permanently into a non-reactive, solid matrix. Furthermore, the upregulation of tyrosine metabolism also sustains the melanisation pathway, providing direct systemic antioxidant defence (Abbasi, 2025; Janssen *et al*., 2019).

The cellular management of iron absorption within the midgut epithelium constitutes the next critical level of protection against systemic iron toxicity. In this study, *A. domesticus* was reared on a grain-based chicken feed diet enriched with FeSO_4_. Although explicit iron speciation of the diet was not conducted, the dietary iron pool is expected to be predominantly non-heme, combining native plant-derived ferric iron (existing primarily as Fe^3+^ with minor Fe^2+^ fractions (De Brier *et al*., 2016; Singh *et al*., 2013)) with supplemented ferrous iron (Fe^2+^). While the added FeSO_4_ is intended to provide a direct bioavailable iron source, the Fe^3+^ ions require ferrireductase activity prior to cellular uptake (Hernández-Gallardo *et al*., 2024). Once present within the midgut lumen in this divalent state, these iron ions form a collective Fe^2+^ pool that is transferred into the midgut epithelium by Malvolio (Mvl), the insect ortholog of the mammalian divalent metal transporter 1 (DMT1) (Mandilaras *et al*., 2013). To survive extreme midgut iron exposure, *A. domesticus* may rely on intestinal adaptation mechanisms, that potentially involve downregulating the apical importer Mvl alongside ferritin-mediated sequestration within midgut epithelial cells. These cells might also utilize secretory ferritin pathways to exocytose excess iron back into the lumen for excretion, thereby preventing systemic iron toxicity (Gorman, 2023; Mandilaras *et al*., 2013; Tang and Zhou, 2013). In line with these regulatory concepts, our work proves that the crickets retain a regulated responsiveness to dietary iron manipulation, showing a moderate shift from a modest 21.38 mg Fe/kg DM to 118.34 mg Fe/kg DM.

Beyond iron enrichment, the quantified elemental data revealed that macroelements (Na, K, Ca, Mg, P, S), microelements (Zn, Cu, Co, Se, Mo), and heavy metals (Cd, Pb, As) remained remarkably stable, regardless by the iron feeding level, with only Mn showing significative differences. This indicates that the high-dose iron feeding did not induce generalized systemic toxicity or disrupt epithelial membrane integrity within the digestive tract, which would typically trigger a leakage or deregulation of essential electrolyte pools. Instead, these findings suggest an underlying interrelation between iron levels and trace element transportation and homeostasis inside the crickets. The significant reduction in manganese between the CTR and the Fe_100mM groups is likely a consequence of competition with iron in the use of the specific homeostatic systems. Indeed, in biological systems, manganese primarily exists as a divalent cation (Mn^2+^) and therefore shares the identical valence state of the luminal Fe^2+^ pool generated after the ferrireductase activity. Consequently, both cations compete for the same apical midgut epithelium entry transporter. The downregulation of Mvl1 under high iron load (Missirlis and Oliveira, 2022) likely amplifies this competitive antagonism. Furthermore, although their exact presence in *A. domesticus* remains to be fully characterized, putative homologs of the zinc-regulated transporters ZIP8 and ZIP14 (SLC39A family) may also contribute to this shared absorption route (Winslow *et al*., 2020). Consequently, an extreme dietary overload of iron could competitively inhibit manganese uptake across these overlapping multi-transporter gateways. In *Drosophila melanogaster*, competitive inhibition has been reported between iron and manganese, leading to tissue manganese depletion at even sufficient dietary supply (Pankau and Cooper, 2022). This suggests evolutionary conservation of metal ion transport routes and homeostasis across insect taxa (Pankau and Cooper, 2022). This strong Fe-Mn competition may have negative implications for cricket farming and biofortification strategy by compromising important physiological status and reduced nutritional value of crickets for human consumption.

Interestingly, although not significant, zinc level in insects increased when low iron levels were present in diet, it peaked in the Fe_10mM crickets and then decreased. Iron-zinc competitive inhibition is well-documented in vertebrate nutrition, in which supplemental iron can inhibit zinc absorption (Fairweather-Tait, 1995; Sandstrom *et al*., 1985; Troost *et al*., 2003). However, there are recent suggestions that similar competitive interactions occur in other insects such as *T. molitor* and *H. illucens* (First *et al*., 2024). Therefore, if such an interaction exists, zinc should be monitored for effective biofortification in crickets as well. A similar decline trend was also observed for cadmium under iron treatment. Because insects share intestinal pathways for essential trace elements and non-essential heavy metals (Allard *et al*., 2026; Keil *et al*., 2020), this decline strongly points toward a competitive exclusion mechanism during epithelial transport as already suggested by Vesey (2010) in vertebrate animals. Like manganese, cadmium likely shares the Mvl transporter (Allard *et al*., 2026). Under extreme iron overload, this carrier might become saturated by Fe^2+^ ions, outcompeting cadmium and explaining its observed reduction. Investigating this iron-metal crosstalk within *A. domesticus* and other mass-reared species will expand our understanding of insect metal regulation, providing a clearer picture of the true physiological complexity of the metal homeostasis in insects produced for food and feed. Dietary iron also appears to be responsible for promoting a progressive shift in the cricket’s amino-acid profile. A prime example is phenylalanine hydroxylase (PAH), a strictly iron-dependent enzyme (Du and Zhou, 2025). The enrichment of tyrosine peaking at Fe_25mM suggests maximized PAH activity. However, the simultaneous accumulation of both the substrate phenylalanine and the PAH reaction product tyrosin indicates that their downstream turnover or degradation pathways may be saturated or limited at this concentration. This enzymatic bottleneck could point toward a potential preservation strategy, where *A. domesticus* limits the catabolism of these aromatic amino acids to potentially retain them for structural or protective demands, such as cuticle sclerotization and immune defense (Vavricka *et al*., 2014). The structural necessity of this pathway has recently been demonstrated in *Gryllus bimaculatus*, where disruption of tyrosine metabolism directly impaired chitin matrix deposition, leading to defective cuticle formation (Bai *et al*., 2026). Beyond that, the PCA biplot confirmed iron-supplemented crickets, characterized by an enrichment of core metabolic amino acids— including the TCA-derived aspartic and glutamic acids, alongside alanine, proline, serine, and threonine—reflecting a coordinated upregulation of systemic nitrogen and energy pathways. Thus, iron supplementation at Fe_25mM may support cricket’s capacity to convert dietary nitrogen into amino acids, reducing nitrogenous waste and increasing protein deposition (Ren *et al*., 2026). This is consistent with the higher feed conversion ratios we observed at moderate to high levels of iron supplementation. However, the intermediary clustering of Fe_100mM iron supplementation suggests that metabolic stress may occur at higher iron supplementation in crickets as high levels of iron generate reactive oxygen species via Fenton chemistry that damages amino acid side chains, especially the sulphur-containing methionine and cysteine.

## Conclusion

This study confirmed that iron biofortification can affect edible crickets, in particular, altering the mineral composition, amino-acid profile, and rearing performances. Our findings confirmed that dietary iron supplementation within the range of 10-25 mmol Fe does not affect survival, growth, nutritional parameters, and microbiome composition of edible crickets. We did observe a Fe-Mn interaction at moderate iron enrichment and macronutrient stability, suggesting that crickets can manage trace element balance when faced with minor dietary challenges. These results are incredibly promising, suggesting that crickets are able to tolerate relatively high levels of iron supplementation without risk to insect or consumer health. These results are particularly encouraging in the context of producing protein biofortified with iron in areas of the global south where ID and IDA are particularly prolific problems and where eating insect protein is accepted.

Some development steps must now be taken in order to establish biofortified crickets as a safe iron source. While a short-term tolerance to altered Fe-Mn and Fe-Zn ratios under controlled laboratory conditions needs to be considered, the exact minimal requirements for these minor components for crickets under industrial rearing conditions remain undefined. Crucially, subtle physiological impairments or the outcome of marginal zinc and manganese deficiencies may only manifest under commercial rearing stressors or pathogen exposure. Concurrently, profiling host transcriptomics and tissue-specific transport pathways will be necessary in order to clarify how crickets functionally regulate iron uptake under iron supplementation. Integrating these host responses with microbiome-derived metabolic shifts will help determine whether the microbial community actively sequesters iron or enhances its chemical solubility within the gut lumen. Ultimately, incorporating robust bioavailability assessments in future studies—utilizing either in vitro digestion models coupled with intestinal cell lines (such as Caco-2) or in vivo consumer trials—is crucial to verify the extent to which the iron within the insect matrix is effectively bioavailable, providing an evidence-based strategy to enhance the nutritional value of edible insects.

## Acknowledges

Author G.R. would like to thank Mrs. Eleni Gousgoula, Dr. Adriana Poccia and Mrs. Gabrielle Wegner for supporting the chemical analysis. M.M. and C.S. thank Gabriele Pohl for her assistance in the metal analysis. HH thanks the Foundation for Food Safety and Consumer Protection (Stiftung für Lebensmittelsicherheit und Verbraucherschutz) for their generous financial support.

