## Supplementary figures and images for "Impact of iron-enriched diets on the nutritional composition and microbiome diversity of house crickets (*Acheta domesticus*)"

### Figure S1

**Supplementary Figure S1.** Overview of the crickets rearing procedure and experimental set-up.

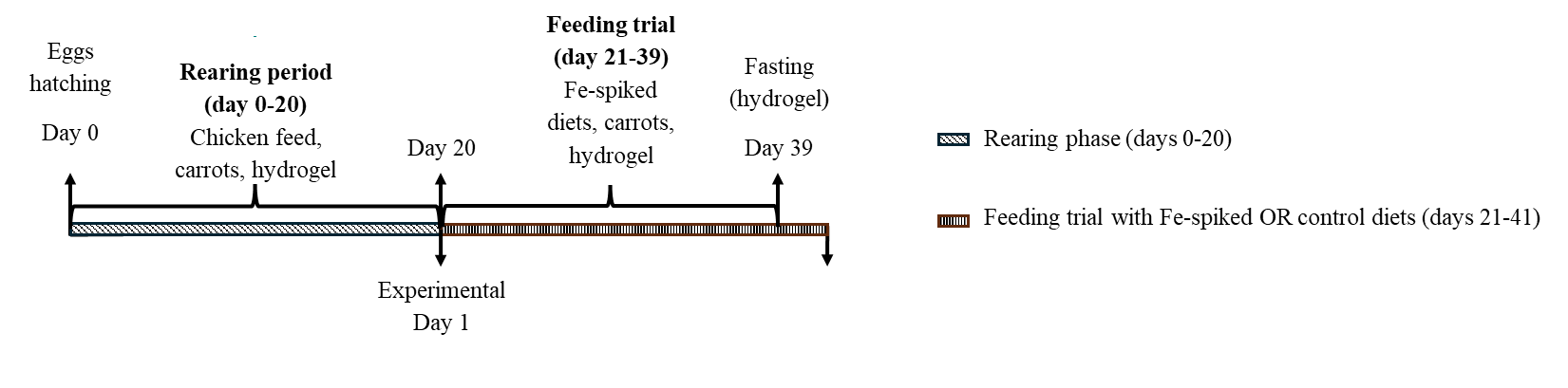
