## Supplementary material for "Impact of iron-enriched diets on the nutritional composition and microbiome diversity of house crickets (*Acheta domesticus*)": Table S1

**Supplementary Table S1.** Recovery rate (average ± standard deviation) and quantification limit of the metals measured by the ICP-MS/MS.

| Element | Collision gas | Mass shift | Recovery rate (%) | Limit of quantification (µg/L) |
| --- | --- | --- | --- | --- |
| Na | He | 23 🡪 23 | 102.6 ± 0.2 | 17.9 |
| Mg | He | 24 🡪 24 | 109.6 ± 3.4 | 0.9 |
| P | O_2_ | 31 🡪 47 | - | 2.5 |
| S | O_2_ | 32 🡪 48 | - | 7.4 |
| K | He | 39 🡪 39 | 102.4 ± 4.4 | 29.5 |
| Ca | He | 44 🡪 44 | 108.7 ± 12.3 | 27.5 |
| Mn | He | 55 🡪 55 | 95.1 ± 9.3 | 0.04 |
| Fe | He | 56 🡪 56 | 92.6 ± 6.4 | 0.50 |
| Co | He | 59 🡪 59 | 85.8 ± 7.9 | 0.005 |
| Ni | He | 60 🡪 60 | - | 0.17 |
| Cu | He | 63 🡪 63 | 95.1 ± 8.0 | 0.05 |
| Zn | He | 66 🡪 66 | 94.1 ±11.0 | 1.28 |
| As | O_2_ | 75 🡪 91 | 91.2 ± 9.7 | 0.01 |
| Se | O_2_ | 80 🡪 96 | 84.0 ± 6.0 | 0.11 |
| Mo | He | 95 🡪 95 | - | 0.01 |
| Cd | He | 111 🡪 111 | 97.1 ± 16.2 | 0.01 |
| Pb | He | 208 🡪 208 | 91.1 ± 18.2 | 0.03 |
| Ge | He, O_2_ | 72 🡪 72 | - | - |
| Rh | He, O_2_ | 103 🡪 103 | - | - |
