## Supplementary material for "Impact of iron-enriched diets on the nutritional composition and microbiome diversity of house crickets (*Acheta domesticus*)": Table S2

**Supplementary Table S2.** Experimental conditions for metal quantification via ICP-MS/MS.

| Plasma forward power | 1550 W |
| --- | --- |
| Plasma gas | 15.0 mL/min |
| Auxiliary gas flow | 0.90 mL/min |
| Nebulizer gas flow | 0.99 mL/min |
| Nebulizer type | MicroMist |
| Collision gas | 4.5 mL/min (He), 30 % (O2) |
| Calibration range | 1.0 – 5000 (Na, Mg, K, S, P)  2.0 – 10000 (Ca)  0.2 – 1000 (Fe, Cu, Zn)  0.01 – 50 µg/L (Mn, Co, Ni, As, Se, Mo, Cd, Pb) |
