## Supplementary material for "Impact of iron-enriched diets on the nutritional composition and microbiome diversity of house crickets (*Acheta domesticus*)": Table S3

**Supplementary Table S3.** Amino acid profile of *A. domesticus* nymphs reared on diets supplemented with different levels of iron. Values are presented as mean ± SE (n = 5). Different letters within the same row indicate statistically significant differences (one-way ANOVA followed by Tukey's HSD post hoc test; P < 0.05).

|  | CTR | Fe_1mM | Fe_5mM | Fe_10mM | Fe_25mM | Fe_100mM |
| --- | --- | --- | --- | --- | --- | --- |
| Ammonia (mg/g DM) | 4.86 ± 0.39 d | 4.32 ± 0.16 d | 5.54 ± 0.39 cd | 6.34 ± 0.38 bc | 7.84 ± 0.16 a | 7.42 ± 0.31 ab |
| Aspartic Acid (mg/g DM) | 30.60 ± 3.17 c | 27.38 ± 1.17 c | 32.86 ± 1.88 bc | 38.11 ± 1.41 ab | 44.24 ± 0.94 a | 42.66 ± 1.34 a |
| Threonine (mg/g DM) | 14.53 ± 1.11 c | 12.79 ± 0.48 c | 16.34 ± 1.14 bc | 19.00 ± 0.79 ab | 21.90 ± 0.66 a | 21.11 ± 0.58 a |
| Serine (mg/g DM) | 13.97 ± 1.61 c | 12.15 ± 0.57 c | 16.89 ± 1.63 bc | 19.26 ± 0.99 ab | 22.85 ± 0.40 a | 21.69 ± 0.60 a |
| Glutamic Acid (mg/g DM) | 42.65 ± 4.94 bc | 38.53 ± 1.57 c | 47.82 ± 3.16 bc | 55.06 ± 1.96 ab | 66.21 ± 2.31 a | 62.37 ± 2.36 a |
| Proline (mg/g DM) | 28.04 ± 3.36 a | 21.76 ± 1.35 a | 29.28 ± 2.09 a | 30.34 ± 1.71 a | 26.83 ± 1.56 a | 30.03 ± 1.40 a |
| Glycine (mg/g DM) | 18.38 ± 1.62 ab | 15.75 ± 0.77 b | 19.97 ± 1.28 ab | 16.63 ± 4.19 b | 3.18 ± 3.18 c | 28.54 ± 1.31 a |
| Alanine (mg/g DM) | 35.79 ± 2.85 cd | 30.91 ± 1.68 d | 39.75 ± 2.66 cd | 43.60 ± 3.11 bc | 57.24 ± 2.05 a | 54.41 ± 2.50 ab |
| Valine (mg/g DM) | 18.06 ± 1.59 bc | 14.28 ± 0.89 c | 19.78 ± 0.95 bc | 21.46 ± 2.49 b | 31.84 ± 1.15 a | 28.84 ± 1.20 a |
| Methionine (mg/g DM) | 5.33 ± 0.51 c | 4.60 ± 0.24 c | 5.53 ± 0.40 c | 7.37 ± 0.36 b | 9.18 ± 0.39 a | 7.76 ± 0.29 ab |
| Isoleucine (mg/g DM) | 12.98 ± 0.89 cd | 11.26 ± 0.45 d | 13.26 ± 0.71 cd | 15.56 ± 0.74 bc | 19.18 ± 0.63 a | 17.13 ± 0.72 ab |
| Leucine (mg/g DM) | 29.10 ± 2.55 bc | 25.13 ± 1.01 c | 29.61 ± 1.75 bc | 35.22 ± 1.93 ab | 44.17 ± 1.50 a | 38.83 ± 1.49 a |
| Tyrosine (mg/g DM) | 21.59 ± 1.71 d | 16.77 ± 0.36 d | 23.60 ± 1.85 cd | 29.49 ± 2.52 bc | 37.48 ± 1.66 a | 35.87 ± 1.32 ab |
| Phenylalanine (mg/g DM) | 12.16 ± 1.98 c | 10.92 ± 0.72 c | 12.40 ± 0.95 bc | 15.19 ± 1.33 abc | 20.26 ± 1.06 a | 16.86 ± 1.05 ab |
| Histidine (mg/g DM) | 31.97 ± 3.31 bc | 23.75 ± 1.51 c | 30.58 ± 2.70 bc | 40.82 ± 3.24 ab | 51.04 ± 1.57 a | 44.65 ± 2.38 a |
| Lysine (mg/g DM) | 22.81 ± 2.14 d | 20.37 ± 0.73 d | 24.19 ± 1.30 cd | 29.34 ± 1.32 bc | 36.19 ± 1.14 a | 31.54 ± 1.16 ab |
| Arginine (mg/g DM) | 24.79 ± 2.36 c | 20.90 ± 0.75 c | 25.95 ± 1.48 bc | 32.60 ± 1.79 ab | 39.32 ± 1.37 a | 33.45 ± 1.37 a |
