## Supplementary material for "Impact of iron-enriched diets on the nutritional composition and microbiome diversity of house crickets (*Acheta domesticus*)": Table S4

**Supplementary Table S4.** Pairwise PERMANOVA computed on: (a.) Bray-Curtis dissimilarity matrix including starter crickets; (b.) Bray-Curtis dissimilarity matrix excluding starter crickets; (c.) Jaccard dissimilarity matrix including starter crickets; (d.) Jaccard dissimilarity matrix excluding starter crickets.

| a. | | | | | | |
| --- | --- | --- | --- | --- | --- | --- |
| COMPARISON | Df | SumsOfSqs | F.Model | R2 | p.value | p.adjusted |
| 100mmol_Fe vs 10mmol_Fe | 1 | 0.05694001 | 0.540919 | 0.063333 | 0.933 | 0.96 |
| 100mmol_Fe vs 1mmol_Fe | 1 | 0.10046516 | 0.93307 | 0.104451 | 0.572 | 0.96 |
| 100mmol_Fe vs 25mmol_Fe | 1 | 0.07991686 | 0.694602 | 0.079889 | 0.831 | 0.96 |
| 100mmol_Fe vs 5mmol_Fe | 1 | 0.07871394 | 0.690838 | 0.07949 | 0.762 | 0.96 |
| 100mmol_Fe vs Control | 1 | 0.12262422 | 0.986346 | 0.10976 | 0.465 | 0.96 |
| 100mmol_Fe vs Starter_crickets | 1 | 0.34983339 | 3.881247 | 0.392789 | 0.017 | 0.084 |
| 10mmol_Fe vs 1mmol_Fe | 1 | 0.05798516 | 0.63675 | 0.073726 | 0.96 | 0.96 |
| 10mmol_Fe vs 25mmol_Fe | 1 | 0.08431206 | 0.856423 | 0.096701 | 0.546 | 0.96 |
| 10mmol_Fe vs 5mmol_Fe | 1 | 0.06063643 | 0.622983 | 0.072247 | 0.857 | 0.96 |
| 10mmol_Fe vs Control | 1 | 0.07302444 | 0.677945 | 0.078123 | 0.917 | 0.96 |
| 10mmol_Fe vs Starter_crickets | 1 | 0.29054073 | 4.273214 | 0.415957 | 0.018 | 0.084 |
| 1mmol_Fe vs 25mmol_Fe | 1 | 0.10892994 | 1.080084 | 0.118951 | 0.367 | 0.96 |
| 1mmol_Fe vs 5mmol_Fe | 1 | 0.07689351 | 0.770949 | 0.087898 | 0.729 | 0.96 |
| 1mmol_Fe vs Control | 1 | 0.09346802 | 0.848777 | 0.09592 | 0.604 | 0.96 |
| 1mmol_Fe vs Starter_crickets | 1 | 0.29359526 | 4.123547 | 0.407322 | 0.021 | 0.084 |
| 25mmol_Fe vs 5mmol_Fe | 1 | 0.08310824 | 0.775833 | 0.088406 | 0.661 | 0.96 |
| 25mmol_Fe vs Control | 1 | 0.11618497 | 0.988781 | 0.110002 | 0.468 | 0.96 |
| 25mmol_Fe vs Starter_crickets | 1 | 0.31628226 | 3.902647 | 0.394101 | 0.013 | 0.084 |
| 5mmol_Fe vs Control | 1 | 0.08633967 | 0.741821 | 0.084859 | 0.811 | 0.96 |
| 5mmol_Fe vs Starter_crickets | 1 | 0.28225619 | 3.547843 | 0.371586 | 0.024 | 0.084 |
| Control vs Starter_crickets | 1 | 0.36735149 | 3.933108 | 0.395959 | 0.016 | 0.084 |
| b. | | | | | | |
| 100mmol_Fe vs 10mmol_Fe | 1 | 0.05694001 | 0.540919 | 0.063333 | 0.914 | 0.955 |
| 100mmol_Fe vs 1mmol_Fe | 1 | 0.10046516 | 0.93307 | 0.104451 | 0.593 | 0.955 |
| 100mmol_Fe vs 25mmol_Fe | 1 | 0.07991686 | 0.694602 | 0.079889 | 0.855 | 0.955 |
| 100mmol_Fe vs 5mmol_Fe | 1 | 0.07871394 | 0.690838 | 0.07949 | 0.752 | 0.955 |
| 100mmol_Fe vs Control | 1 | 0.12262422 | 0.986346 | 0.10976 | 0.449 | 0.955 |
| 10mmol_Fe vs 1mmol_Fe | 1 | 0.05798516 | 0.63675 | 0.073726 | 0.955 | 0.955 |
| 10mmol_Fe vs 25mmol_Fe | 1 | 0.08431206 | 0.856423 | 0.096701 | 0.541 | 0.955 |
| 10mmol_Fe vs 5mmol_Fe | 1 | 0.06063643 | 0.622983 | 0.072247 | 0.835 | 0.955 |
| 10mmol_Fe vs Control | 1 | 0.07302444 | 0.677945 | 0.078123 | 0.897 | 0.955 |
| 1mmol_Fe vs 25mmol_Fe | 1 | 0.10892994 | 1.080084 | 0.118951 | 0.361 | 0.955 |
| 1mmol_Fe vs 5mmol_Fe | 1 | 0.07689351 | 0.770949 | 0.087898 | 0.715 | 0.955 |
| 1mmol_Fe vs Control | 1 | 0.09346802 | 0.848777 | 0.09592 | 0.625 | 0.955 |
| 25mmol_Fe vs 5mmol_Fe | 1 | 0.08310824 | 0.775833 | 0.088406 | 0.671 | 0.955 |
| 25mmol_Fe vs Control | 1 | 0.11618497 | 0.988781 | 0.110002 | 0.435 | 0.955 |
| 5mmol_Fe vs Control | 1 | 0.08633967 | 0.741821 | 0.084859 | 0.778 | 0.955 |
| c. |  |  |  |  |  |  |
| 100mmol_Fe vs 10mmol_Fe | 1 | 0.1444186 | 0.761328 | 0.086896 | 0.844 | 0.952737 |
| 100mmol_Fe vs 1mmol_Fe | 1 | 0.1956674 | 1.006297 | 0.111733 | 0.498 | 0.952737 |
| 100mmol_Fe vs 25mmol_Fe | 1 | 0.1660165 | 0.81777 | 0.092741 | 0.787 | 0.952737 |
| 100mmol_Fe vs 5mmol_Fe | 1 | 0.1753665 | 0.874989 | 0.09859 | 0.627 | 0.952737 |
| 100mmol_Fe vs Control | 1 | 0.2227575 | 1.036256 | 0.114678 | 0.389 | 0.952737 |
| 100mmol_Fe vs Starter_crickets | 1 | 0.5263796 | 3.243087 | 0.350866 | 0.014 | 0.084 |
| 10mmol_Fe vs 1mmol_Fe | 1 | 0.1314227 | 0.747657 | 0.085469 | 0.962 | 0.962 |
| 10mmol_Fe vs 25mmol_Fe | 1 | 0.1685727 | 0.914428 | 0.102578 | 0.549 | 0.952737 |
| 10mmol_Fe vs 5mmol_Fe | 1 | 0.1324401 | 0.728662 | 0.083479 | 0.862 | 0.952737 |
| 10mmol_Fe vs Control | 1 | 0.1491451 | 0.75978 | 0.086735 | 0.908 | 0.9534 |
| 10mmol_Fe vs Starter_crickets | 1 | 0.4687617 | 3.41107 | 0.362453 | 0.019 | 0.084 |
| 1mmol_Fe vs 25mmol_Fe | 1 | 0.1984603 | 1.049512 | 0.115974 | 0.38 | 0.952737 |
| 1mmol_Fe vs 5mmol_Fe | 1 | 0.1595022 | 0.855204 | 0.096576 | 0.665 | 0.952737 |
| 1mmol_Fe vs Control | 1 | 0.1674652 | 0.832952 | 0.094301 | 0.751 | 0.952737 |
| 1mmol_Fe vs Starter_crickets | 1 | 0.4707364 | 3.274531 | 0.353067 | 0.024 | 0.084 |
| 25mmol_Fe vs 5mmol_Fe | 1 | 0.1652933 | 0.847327 | 0.095772 | 0.666 | 0.952737 |
| 25mmol_Fe vs Control | 1 | 0.1972555 | 0.941021 | 0.105248 | 0.595 | 0.952737 |
| 25mmol_Fe vs Starter_crickets | 1 | 0.4894644 | 3.154146 | 0.344559 | 0.02 | 0.084 |
| 5mmol_Fe vs Control | 1 | 0.1659626 | 0.801641 | 0.091079 | 0.83 | 0.952737 |
| 5mmol_Fe vs Starter_crickets | 1 | 0.4546045 | 2.996178 | 0.33305 | 0.017 | 0.084 |
| Control vs Starter_crickets | 1 | 0.5360385 | 3.132565 | 0.34301 | 0.022 | 0.084 |
| d. |  |  |  |  |  |  |
| 100mmol_Fe vs 10mmol_Fe | 1 | 0.1444186 | 0.761328 | 0.086896 | 0.813 | 0.962 |
| 100mmol_Fe vs 1mmol_Fe | 1 | 0.1956674 | 1.006297 | 0.111733 | 0.523 | 0.962 |
| 100mmol_Fe vs 25mmol_Fe | 1 | 0.1660165 | 0.81777 | 0.092741 | 0.809 | 0.962 |
| 100mmol_Fe vs 5mmol_Fe | 1 | 0.1753665 | 0.874989 | 0.09859 | 0.656 | 0.962 |
| 100mmol_Fe vs Control | 1 | 0.2227575 | 1.036256 | 0.114678 | 0.362 | 0.962 |
| 10mmol_Fe vs 1mmol_Fe | 1 | 0.1314227 | 0.747657 | 0.085469 | 0.962 | 0.962 |
| 10mmol_Fe vs 25mmol_Fe | 1 | 0.1685727 | 0.914428 | 0.102578 | 0.577 | 0.962 |
| 10mmol_Fe vs 5mmol_Fe | 1 | 0.1324401 | 0.728662 | 0.083479 | 0.861 | 0.962 |
| 10mmol_Fe vs Control | 1 | 0.1491451 | 0.75978 | 0.086735 | 0.914 | 0.962 |
| 1mmol_Fe vs 25mmol_Fe | 1 | 0.1984603 | 1.049512 | 0.115974 | 0.375 | 0.962 |
| 1mmol_Fe vs 5mmol_Fe | 1 | 0.1595022 | 0.855204 | 0.096576 | 0.66 | 0.962 |
| 1mmol_Fe vs Control | 1 | 0.1674652 | 0.832952 | 0.094301 | 0.756 | 0.962 |
| 25mmol_Fe vs 5mmol_Fe | 1 | 0.1652933 | 0.847327 | 0.095772 | 0.657 | 0.962 |
| 25mmol_Fe vs Control | 1 | 0.1972555 | 0.941021 | 0.105248 | 0.544 | 0.962 |
| 5mmol_Fe vs Control | 1 | 0.1659626 | 0.801641 | 0.091079 | 0.84 | 0.962 |
